# Gone in 40 years, the curious case of the Himalayan Quail: an attempt at rediscovery and implications for conservation

**DOI:** 10.1101/2020.07.14.201137

**Authors:** Paul Pop, Puneet Pandey, Randeep Singh

## Abstract

Himalayan Quail (*Ophrysia superciliosa*) is a Galliforme species currently classified as Critically Endangered by the IUCN. Given the fact that the species has only been recorded over a period of 40 years in the 19^th^ century, this classification needs more scrutiny. The aim of this study was to explore the possibility of detecting the species in their known historical distribution in the lower Himalayas, and understand if possible, the factors which led to their decline or extinction. The potential habitats in districts of Nainital and Dehradun of Uttrakhand (India), where the species which has been recorded between 1836 to 1876, have been scoured thoroughly insofar as logistics and other factors permitted using trail and point transects. We did not detect any sign of the species in any of the areas, thereby corroborating the results of past surveys/expeditions in search of this species. Domesticated/feral species (grazing pressure and hunting by dogs), natural predation by species such as Yellow-throated Marten (*Martes flavigula*), hunting by humans, population growth and resulting land-use change, biogeochemical events, and tourism may all have contributed in varying degrees to their decline or extinction. It was found that most, if not all, proposals of strategies to find the species have been thus far unemployed or futile. We suggest improving the chances of re-discovering this species using tools such as molecular/genetic analysis and Unmanned Aerial Vehicles. Management recommendations include stress on grazing laws, sterilization of dogs, awareness about unchecked human population growth and measures for limiting the effects of unsustainable tourism.

## 1. Introduction

It is common scientific knowledge that species are going extinct at an astounding rate, especially within vulnerable taxa like Amphibia and Mammalia (Ripple et al., 2017). Although birds are safer compared to the afore-mentioned taxa due to abilities such as flight, they’re also under the hammer of the ongoing ‘Sixth Mass Extinction’ (Ceballos et al, 2017). This is more pronounced for taxa like Galliformes, characterized by ground-dwelling habit. Since the 1500, atleast 150 species of birds have gone Extinct or Possibly Extinct and majority of them were geographically isolated (close to 90%) – they lived in islands (Butchart et al, 2006). The most recently updated IUCN Red List, has logged in 800 species of birds as Vulnerable, 461 as Endangered and 225 as Critically Endangered accounting for a total of 14% of the extant bird species (Tables 1a & 2, IUCN, 2020). According to the latest version of HBW checklist, 78 species of the order Galliformes are threatened (in the Vulnerable, Endangered or Critically Endangered categories) and two are extinct (Table S1). Himalayan Quail aka Mountain Quail (*Ophrysia superciliosa*) (hereafter ‘HQ’) is one such Galliforme species.

### 1.1. Background

Morphology and anatomy for HQ had been briefly described by several authors (example: Murray 1890). Accounts with ecological descriptions have been recorded too (example: Oates, 1898). Most of these works are recycled information from previous works, with little to no new information. These were further compiled and elaborated in the detailed report of Rieger and Walzthöny (1993) – probably the most comprehensive of all accounts of the species till date. But a few minor details have been missed in this report, which can be found in dated manuscripts, such as:

- HQ is Palaearctic in origin (Ripley, 1959).
- This species may be better classified under the subfamily Perdicinae (Ridgway & Friedmann 1946).

#### 1.1.1. Previous surveys

The earliest recorded attempt at searching for the species was by Salim Ali in 1977, followed by Ravi Sankaran from the BNHS between 1987 and 1990 (described as having had surveyed Mussoorie in 1987 with no sightings or indications (Rieger and Walzthöny, 1993)), then by Ingo Rieger and Doris Walzthöny in 1989. This was succeeded by the group of researchers - Rahul Kaul, Tehmina Shafiq and Salim Javed from the World Wide Fund for Nature (WWF)-India, from 1998 to 1999. Uttarakhand Forest Department (via the Nainital Zoo) had initiated a “Mission Himalayan Quail” on October month of 2013, which was supported by WWF-India. These are the recorded attempts in search for the HQ (Banerjee, 2015).

Baral (2009) had suggested an expedition to Nepal’s western region as only one documented study by Dillon S. Ripley in mid 20^th^ century (Ripley, 1950) had been carried out in Nepal (in Rekcha village from 25^th^ December to 2^nd^ January). Baral and colleagues have been reported to have conducted surveys recently (BirdLife International, 2018), but the specifics has not been made available in the public domain. Same goes for updates on surveys carried out in India by Kalsi and colleagues (mentioned in Kalsi et al, 2007).

#### 1.1.2. Claimed sightings

There have been quite a number of unsubstantiated claims of sighting the bird, after the last confirmed record was made (Reiger and Walthonzy, 1993). Some of them have been found to be false due to conflicting accounts and inconsistent narrative. The rest of them (BirdLife International, 2018) remains uncorroborated.

### 1.2. Aim

To explore the possibility of rediscovery of the Critically Endangered HQ in the lower Himalayas and understand the factors/causes which led to their decline or extinction.

### 1.3 Study Area

This study was carried out in the lower Western Himalayan districts of Nainital and Dehradun of Uttarakhand, India (both names referring to districts in the text hereafter), with focus on Nainital town and Mussoorie, where the species had been historically reported (Fig. 1, Plate S1). They are characterised by variation in altitude, as well as weather, with warm and sunny summer, high-rainfall monsoons and very cold winters with snowfall (depending on the elevation). Both Mussoorie and Nainital town are occupied by conifer mountain forests.

**Fig. 1.**
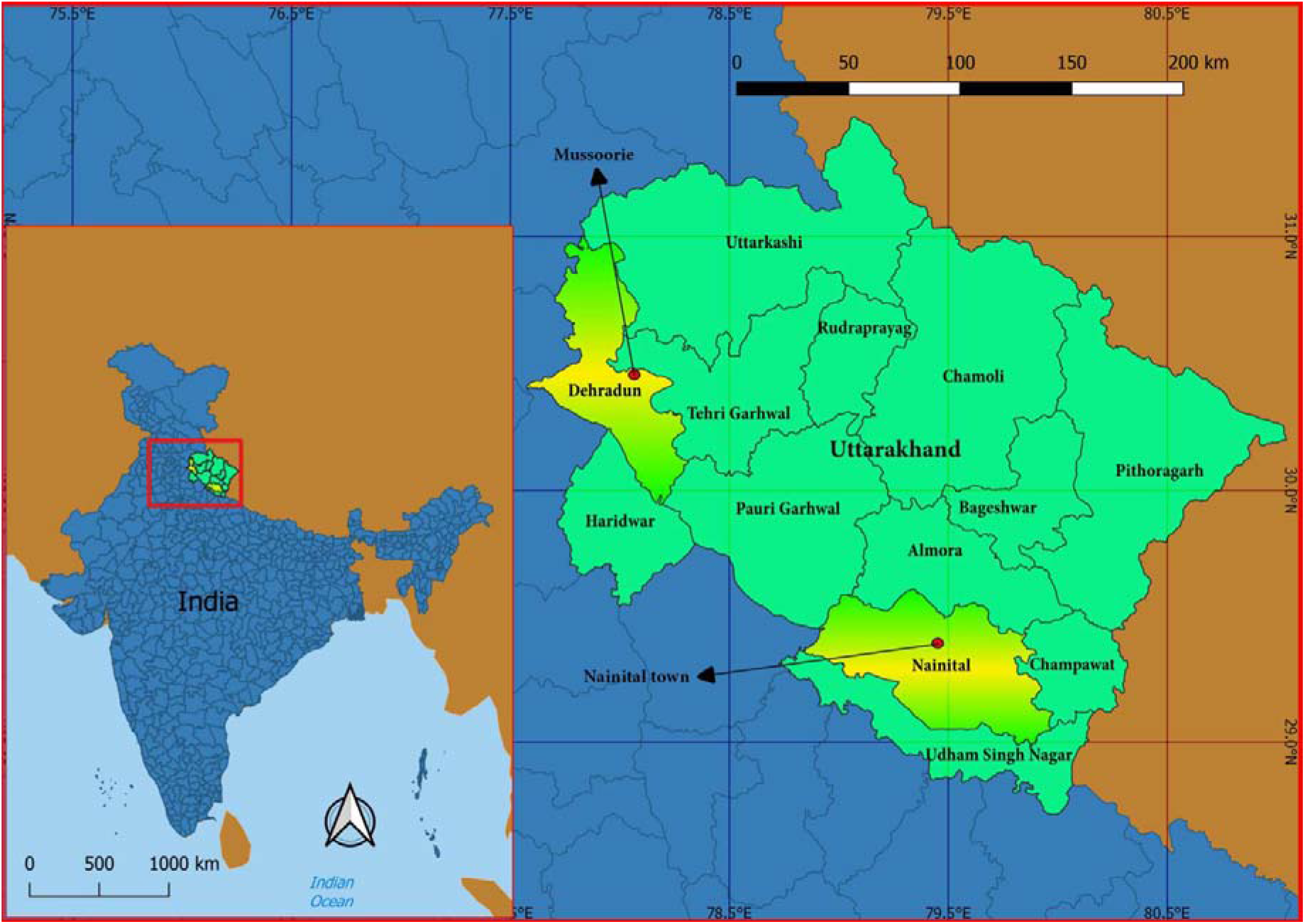
Study Area Map

## 2. Methodology

### 2.1. Site Selection

The sites were decided based on review of literature and later on reconnaissance ground surveys for accessibility and safety. Reiger and Walthonzy (1993), as well as Dunn et al (2015) were used as primary references for the same.

### 2.2. Field Methodology

The investigation was carried out from 20^th^ September to 31^st^ December, 2018 with 65 days spent in the field split between Nainital (45 days) and Dehradun (20 days). Surveys were conducted within selected trail transects and point transects within the trails. The surveys were carried out in different times of the day for the same sites to account for potential temporal change in habits of the species, although such pheasants are generally more active during dawn and dusk. Surveys were carried out using the mud/dirt/gravel trails and small trails made by domesticated and wild bovines (Goral (*Nemorhaedus goral*)) in addition to Barking Deer (*Muntiacus muntjac*) and Indian Wild Boar (*Sus scrofa*). Locations covered were grasslands/brushwoods/Bamboo patches.

Since there are no recordings of the calls of HQ for callbacks surveys, the author followed the method adopted by Reiger and Walthonzy (1993) with some customization to conditions. It involved stealthily navigating into, and through any potential Quail habitat by making the least possible noise, and then silently waiting motionless at some strategic points to detect any avian motion and listen for any unique calls. Memorization of several vocalizations of all birds and mammals usually occurring in the area had been done beforehand, and a digital call edition of a bird guide was installed in phone for additional support. If some bird calls were unidentifiable during the time, it was recorded, and checked later using online repositories.

A lookout was maintained for indications like feathers, typical of the species (droppings and feeding traces as indications are quite difficult to spot under the grasses) as described and illustrated in Reiger and Walthonzy (1993). Areas far away were scanned thoroughly and slowly through a pair of binoculars. Notes were diligently kept. Google maps was used for navigation, GPS for recording important coordinates and a compass for recording slopes.

## 3. Results

After extensive surveys of potential habitats in Nainital and Dehradun, no signs of HQ had been detected in any of the areas surveyed (approximately 6.5 Km^2^ in Nainital and 3.1 Km^2^ in Dehradun). There are very few potential habitats left in the areas surveyed, as most of the historical range has been turned into agricultural land or used for tourism purposes. Those which still exist, is heavily to moderately disturbed by anthropogenic causes. Most of the slopes with large patches of long grasses were found to lie in the south-facing direction, which is in concordance with the observation of Reiger and Walthonzy (1993). In both districts, there is abundance of insects, and in some places, berries, both of which probably formed a part of the species’ diet, in addition to seed grasses (Oates, 1898) (Plate S2).

### 3.1. Nainital

Focus in Nainital town was at Sher ka danda, the place where the last verified record of the species had been made in 1876 (Rieger & Walzthöny, 1993). During the stay in Nainital town, the area had been intensively surveyed. In Nainital, other areas surveyed by Reiger and Walthonzy (1993) had also been surveyed – Sattal (twice) and Bhimtal (once). In addition, Cheer Nature Trail (7 times), other parts of Nanda Devi Bird Conservation Reserve (twice), Pangot (twice) and China Point (once) had been surveyed. Other areas include the Kahal Jungle routes (7 times), Tiffin Top Route (8 times), grassy slopes close to Neelkanth Hotel (twice) and the routes to, and surrounding the peak adjoining Naina/China Peak (11 times).

Surveys carried out in Nainital showed that most of potential habitat is greatly disturbed by grass-cutting activity (for fuel and fodder) and grazing. Tourist and locals also disturbed the habitat by drinking, smoking and partying in the area, the after-effect of which is rampant littering. The tourists and local students were observed to bring portable speakers while trekking and play the music at high(est) volume, creating noise pollution in the area. Such locations were mostly devoid of avifaunal activities.

The peak next to China Peak was the first to be explored (hereinafter, “Boar Peak”). The south-facing slope of Boar Peak on the side away from the climbing route is grassy, but the height of the grasses is quite low, due to the regular grazing and grass-cutting happening in the area. Further along the ridge situated next to this peak, has grassy expanses but was found to have higher grass-collection impact than the one close to the peak. The backside of the peak didn’t have much grassland, so lesser number of surveys was dedicated to it.

The area surrounding Kahal Jungle had some patches of grasslands, but not very contiguous and had buildings adjoining them. Tiffin Top routes surveyed had some grassland patches, but the heavy presence of raptors such as Steppe Eagle (*Aquila nipalensis*) and Himalayan Griffon (*Gyps himalayensis*) makes Pheasant inhabitation unlikely. This part of the Ayarpatta also had a good amount of bamboo habitat used by Kalij Pheasant (*Lophura leucomelanos*) and Hill Partridge (*Arborophila torqueola*), which could also host HQ, although historical records don’t indicate this. Additionally, the contiguous grassy slopes close to Neelkanth Hotel were surveyed. They were heavily impacted by grazing.

China Peak, the highest peak in the area at 2600 m, opposite to the Boar Peak was surveyed repeatedly. Cheer Pheasant (*Catreus wallichi*) was sighted here several times, not flushing until the observer was less than 1 m away, a behaviour also recorded to be typical of HQ. The peak was defined by a steep cliff-face with grasses, which was recorded as a typical habitat of HQ. Given these two facts, a habitat ideal for Cheer Pheasant seems to be ideal for HQ too. Grassland patches on the same ridge as China and seen from Boar Peak were also visited.

In addition, the grassy slopes en-route to Pangot was surveyed twice, while travelling via the Kilbury road. The ‘Cheer Point’ (different from Cheer Nature Trail) – a very expansive grassland area, found 10-15 Km after the Ghuggu Khan, was also surveyed. Not much of Sattal and Bhimtal (as well as adjacent Naukuchiytal) had any grassland and is located in unsuitable latitude range. See Plates S3-S6 for images.

### 3.2. Dehradun

In Dehradun, Mussoorie was extensively surveyed. The areas mentioned by Reiger and Walthonzy (1993) that had been surveyed were Camel’s Back Road (twice), Kempty Falls (or places along the Kempty route) (5 times); Cloud Ends and other parts of Benog Wildlife Sanctuary (3 times). In addition, suitable habitats in and on the way to George Everest (4 times) and Dhanaulti (4 times) (including Jabarkhet Nature Reserve, Masrana, Suwakholi, Burans Khanda) have been surveyed.

The same problems that plagues Nainital town also affects Mussoorie and adjoining hill stations, but this area had lesser contiguous patches and probably more disturbance. Camel’s Back and Landour regions are impacted by tourist and military presence and seemed devoid of any activities by Pheasants. The surveys done at George Everest were quite fruitless. It was discovered here that the play of sunlight on the rocky crevices creates figures that resembles the Quail (from a distance 50 m or more). So, future reportings could be scrutinized using the time of the day and slope observed (to calculate angle of sunlight). HQ presumably has better camouflage than many of the cryptically plumaged birds present here. So, when they remain very still, it may become extremely difficult to differentiate them from the background (Plate S7-d).

The area closer to the Doon Valley opposite of George Everest peak seems to have been mined (there was a step-like pattern on the slope). Near the George Everest, the ridges atop the slopes (private area) were used as camping area and fires were lit on the ground. This eliminates the possibility of the species existing on the ridges (atleast the area being used). High grazing intensity (>30 goats in a herd and a few cows) was seen near the peaks, where some patches of long grasses exist. The tourist intensity for the area was high (≥30 people were present during every visit).

During all visits to Kempty region, moderate to high intensity of grazing by goats were observed, resulting in short grasses in this slope. No Galliforme activity was observed during the one time climb to the top (during other visits, the hills were scanned from vantage points), probably because it was just afternoon when the slopes were traversed. When asked whether he had seen the species by displaying illustrations, a local goat-herder responded that the bird has been sighted in the morning, most likely confusing them with another Pheasant species.

The route between Mussoorie and Dhanolti had a lot of grassy slopes which would make excellent habitat for the species, least disturbed by anthropogenic causes. But sightings of numerous raptor species may indicate a high level of natural predation here. Masrana village and adjacent areas along this route was surveyed by climbing the hills located there. There was presence of sheeps in the area (including the grassy slopes), as a woolen textile industry was located in this place. Suwakholi, located along this route, is where one of the claimed sightings (in 1984) had taken place (BirdLife International, 2018). This spot was passed on the way to Dhanolti without any sightings, but the time of the day was not conducive for Quail sighting and the habitat looks healthier than elsewhere. There is a possibility that the claim is true. In Dhanolti, climbing and checking all the grassy peaks was not possible, so scanning using binoculars was done from a) the roadside at Kaddu Khal and b) a farmland close to the peak of Eco Park 1 (a local in this area had knowledge about HQ, but only superficially). See Plates S7-S8 for images.

## 4. Discussion

### 4.1 Possible causes of decline/extinction

#### 4.1.1 Domesticated Species

In India, approximately 40 per cent of land is used as grazing grounds, with around 50% of domesticated animals depending on grazing in forests, grasslands and other such ecologically-sensitive areas in the country as part of the grazing-based livestock husbandry (Singh et al, n.d.). In Uttarakhand, over 70% of land area is grazed. Very few Protected Areas (PAs) have grasslands and nearly 40% of PAs suffer from ‘livestock’ grazing and fodder extraction (Singh et al, n.d.). According to Vanik (2008), grazing is one of the top national level threats to grassland ecosystems in India.

It was reported that grazing has resulted in habitat degeneration and loss in the surveyed habitats of Double-banded Courser (*Rhinoptilus bitorquatus*) (Bushan, 1986) and Forest Owlet (*Athene (Heteroglaux) blewitti*) (Jathar et al, 2015, Mehta et al, 2008). Both these cases illustrate the serious threat posed by grazing to Critically Endangered ‘Lazarus’ bird species in India. Other bird species classified as Critically Endangered in India such as Bengal Florican (*Houbaropsis bengalensis*) are also heavily impacted by grazing (Baral et al, 2012). Grazing pressures in potential HQ habitat is generally high, as described earlier (Plate S9).

It has been reported that domestic or feral dogs (*Canis familiaris*) have been partly the reason for the decline and extinction of six species of birds throughout the world, of which fivehad limited or no flying behaviour (like HQ) (Doherty et al, 2017) (Table S2). Another 72 avian species have dogs as a known or potential threat, of which 12 are Galliformes, among which Cheer Pheasant shares geographic range with that of historical range of HQ (Doherty et al, 2017) (Table S3).

Throughout surveys in Nainital and Dehradun, dogs have been seen close to, or inside potential HQ habitats. In Dehradun, a feral dog was observed to scare away and then chase for a little while, a covey of wild pheasants. But such activity was not observed in or near more isolated places with potential presence of Leopard (*Panthera pardus*) probably because dogs are now a major prey for the former in human-use landscape of Uttarakhand like Nainital (Ahmed, 2010). This problem is accentuated by villagers purposefully keeping dogs near wilderness areas in Uttarakhand to protect the ‘livestock’ (Kala & Kothari, 2013). Although free-ranging cats are a major cause of mortality throughout the world, our study revealed no presence of domesticated/feral cats (*Felis catus),nor* rats, in the suitable habitats of the Quail. So, rats and cats probably had little to no role in the decline and/or extinction of HQ.

#### 4.1.2. Natural Predation

Natural predation refers to predation pressure that would have existed without anthropogenic causes (direct hunting/poaching by humans and predation by introduced domesticated animals). Yellow-throated Marten (*Martes flavigula*), a species of Mustellidae family found throughout the altitudinal range (Mudappa, 2013) of HQ, is known to predate upon birds including Phasianidae members (Parr & Duckworth, 2007). During the study period, several sightings of this Mustellid on 7 different dates were made on all major locations surveyed around Nainital town, of which majority were in groups of 3 (n=5) and the majority were in the month of November (n=4) (Plate S10). The first observation of group strength is probably indicative of their hunting efficiency as more numbers of this agile and stealthy species is likely to increase the damage/mortality rate of their target species. The second observation of the increased sighting (assumed to be indicative of their activities, including hunting) during November (winter) is correlated with the seeding of grasses, during which HQ individuals were recorded most (Reiger and Walthonzy, 1993). These two facts lead to the conclusion that Marten may have had a role in the decline and/or extinction of HQ.

#### 4.1.3. Hunting

Six species of Pheasants have been recorded as being hunted in Uttarakhand (Kaul et al, 2004) of which three were sighted in the study duration: Kalij Pheasant, Hill Partridge, Cheer Pheasant – in the decreasing order of sightings (Plate S11). Although records of hunting for the study sites were not available, a villager informed about certain students engaged in hunting in the Tiffin Top area. As recorded by the Forest Department (FD) and others, local hunting certainly does happen in and around the region, sometimes un-detected by the FD. Since species like Kalij Pheasant and Hill Partridge are easier to spot (especially during dawn and dusk), and their abundance is high in the area, they are more easy targets.

#### 4.1.4. Human Population & Landuse Change

Although the population factor encompasses most of the ones described above, this requires a special section due to its significance as the root of most of most problems. Both Nainital town and Mussoorie have seen tremendous increase in numbers since the time they were first established: ~427 times increase in population within 172 years in the former and ~336 times increase within 184 years in the latter, implying an higher overall growth rate for Nainital town compared to Mussoorie (Fig. S1). If this trend continues, then by the beginning of the next decade (by 2031), the populations in the former and the latter would be ~64,252 and ~48,701, respectively.

The land-use change has been very visible. There has been a marked conversion of grasslands into agricultural farmlands in both Nainital and Dehradun (Plates S12-S13). During this investigation, many terrace farms were witnessed to be occupying the highest peaks in the study areas, which were historically a habitat used by the species (Rieger & Walzthöny, 1993). There has been considerable decrease in the forest cover in most of the hill stations: 50.46 Km^2^ (1960) to 32.72 Km^2^ (1985) – a decrease of 27.6% from the total geographic area of the Mussoorie Muncipal Corporation (64.25 Km^2^) (Pant & Kharkwal, 1997).

#### 4.1.5. Bio-geo-chemical Processes

This factor can’t be discounted, since the mountains are constantly in motion, and as a result, there are regular landslides and erosions. During the course of the study, several minor events were recorded in both Nainital town and Mussoorie. Several major disasters of landslides and floods have been recorded in the area (Eapen, 2014). This is also a cause of temporary or permanent habitat damage (Plate S14).

#### 4.1.6. Tourism

Tourism is a bane for the wildlife, even though it generates some revenue for the Forest Dept., utilized for maintenance of Protected Areas nearby. As described before, there is a lot of noise pollution and soil pollution in the wilderness habitats (Plates S15-S16). Noise pollution is also generated by the constant movement of vehicles. The number of tourists visiting Nainital was 4.71 lakh during 1983 and in ’88, it jumped to 5.6 lakh (Joshi & Pant, 1990).The number had gone down to 0.89 lakh by 2009 and later rose to 9.34 lakh by 2018 (Uttarakhand Tourism Board, personal communication, February 18, 2019) (Figure S2). Over three and a half decades (1983-2018), the numbers have gone up by almost twice (98.3% increase).

The number of domestic tourists in both Nainital town and Mussoorie has always constituted the bulk, and in the years 2009-2018, they formed about 99.99% of all visitors (and presumably, it’s the same or similar proportion in the preceding years as well). So, it’s no surprise that littering and nuisance in the wilderness areas are high. As observed, domestic tourists are more blasé towards the environmental degradation in India. The influx of visitors is so bad in Nainital that the local police once had to put up banners saying “housefull” to deter more vehicles from entering the hill station.The effects of tourism may not have been directly felt by the Quails, but could have had a part in their decline and/or extinction mostly via indirect consequences.

### 4.2. What’s the verdict?

Bermuda Petral/Cahow (*Pterodroma cahow*), an island nesting seabird, had been rediscovered after 285 years of no records in 1916 where their habitat was very heavily damaged, in addition to the direct cause of mortality by introduced domesticated species by sailors (Madeiros, 2012). The threats faced are similar in the case of HQ, and their habitat can be considered as ‘sky islands’ (Warshall, 1994). So, there exists a possibility (however small) that the Quail still survives and may be rediscovered, since they have not been re-sighted only within half that gap period - that of 143 years.

‘Lazarus species’ is a term originally used for species re-appearing after gaps in paleontological records (Flessa & Jablonski, 1983), and later co-opted/extended to include rediscovery of species in recent times, considered extinct in paleontological records (Dawson et al, 2006) or in recent natural history records (Meijaard & Nijman, 2014). When it comes to ‘Lazarus’ species of birds in India, thought to be extinct and rediscovered after decades, there are three prominent examples in recorded history – one of Double-banded Courser, rediscovered in 1986 after 86 years of no confirmed record (Bhushan 1986), and second that of Forest Owlet, rediscovered in 1997, 113 years after a verified record (King & Rasmussen, 1997). The third example - Manipuri Bush-quail (*Perdicula manipurensis*) (Pitches (Comp.), 2006) - is even more significant, not because of the rediscovery interval (it’s lower than the other two – 70 years), but of the similarity of one particular behaviour between this species and HQ - that of skulking (Oates, 1898).

World Conservation Monitoring Centre (1990, 1999) and Norton et al (1990) declared/considered HQ extinct in India without a shadow of doubt (but with a clause of “under review”, presumably for other parts of their range – Nepal). Newton (1998) had even given an approximate date of extinction for this species – 1868, eight years before the last verified record! But, such condemnation is scientifically uncalled for (and in the second case, outright fallacious), given the previous examples. Fifty years was the arbitrarily chosen rediscovery period chosen by IUCN in earlier times. The Red List in 1988 (The IUCN Conservation Monitoring Centre) did not list them as extinct, but stated that the taxon is under review. IUCN declared them extinct in the 1994 Red List (Groombridge (Ed.), 1993) and later on, changed their status as Critically Endangered which remains the same to this date. It is interesting to note that eBird denotes the species as extinct.

Researchers have mostly described the species to be “possibly extinct” or “maybe surviving” and similar descriptors indicating uncertainty. There exist no scientific consensus on the subject, and there should not be, since no consensus should be made in the absence of data. Any statistic based on tendencies for survival or extinction, as reported by scientists cannot be used as a verdict, but merely an indicator. Butchart et al (2006) have classified them as “not considered to be Probably Extinct” with “evidence for extinction” as habitat loss and the “evidence against” as unconfirmed reports, even though they claim that sufficient surveys have been carried out, and the habitat remains (all these claims being highly questionable).

Although declaration of HQ as extinct maybe justified from a scientific perspective, it can be a blow from a conservation perspective. The possible presence of the pheasant species is one strong argument that can be used to preserve the habitats in their historical range. If this is gone, land-use change for human and human-related activities (a lot of potential habitat is under privately-owned estates) will result in adding more to habitat loss and threats to vulnerable species.

### 4.3. Possible Future Directions

Before giving our own suggestions/recommendations, let’s look at the ones given by authors before.

- Informing locals about the Quail (Rieger & Walzthöny, 1993).

Comment: After interaction with a number of local people in Nainital town and Mussoorie, it is clear that none of them have a clue about the species, nor their significance (even well educated people). Pamphlets have been reported to have been used by BHNS in the past. WWF-India has stated that they have given out pamphlets on behalf of the Uttarakhand Forest Dept. on a very recent endeavour to search for the species. But, whether the pamphlets were translated to local languages (like Hindi), necessary for many who are unable to read English, was not mentioned.

- Announcing a reward for proof of sighting of this species (Rieger & Walzthöny, 1993).

Comment: Uttarakhand Forest Department had already announced a huge sum of Rs 1 Lakh for anyone who has “irrefutable evidence of Himalayan Quail’s presence”. Nobody has claimed this reward so far. Although the reward is a very good idea, there is a major problem with getting “irrefutable evidence”. Irrefutable proof is likely be photograph(s), dead specimen(s) (unlikely that this will be reported, since it may be perceived as result of hunting by the individual(s)) and feather(s) (also unlikely, as the level of knowledge required for its proper identification by locals are high, in addition to the fact that such collections are criminalized under WPA, 1972). A solution might to be providing a suitable camera to each small villages living near to, or inside the wilderness areas.

- Using a trained dog for seeking out the Quails (Rieger & Walzthöny, 1993).

Comment: Although trained/disciplined dogs regularly visit the trails by morning walkers and joggers, none of them have been reported to have sensed or seeked out any Quails. Close to Burans Khanda, an observation of a dog chasing away a covey of Pheasants was found. So, dogs certainly seek out birds (especially ground birds like Pheasants). If highly trained dogs are brought along for the surveys, they may be able to locate and direct humans to sightings, but the terrain is often too precipitous for dogs.

- Listening for the vocalizations of HQ (Rieger & Walzthöny, 1993).

Comment: This is somewhat pointless, as has been revealed by the current study. This is because there is no recording of the vocalizations for the species and several birds vocalize in the same/a similar manner as has been described in literature. No unusual vocalizations, which could be that of HQ were heard during the study.

- Reducing the numbers within the searching party (Rieger & Walzthöny, 1993).

Comment: The smaller the number of surveyors, the larger the chances of bird sightings is an axiom, largely accepted as true. In the current study, only one person had surveyed the area, thereby increasing chances to maximum. Problem here is the need to lookout for dangers of wild animal attacks while alone.

- Grass burning for flushing the Quails (Rieger & Walzthöny, 1993).

Comment: Rieger & Walzthöny had condemned the use of grass-burning just for flushing out the species, but condones this technique as long as it is common. We condemn grass-burning in both these circumstances, as human-started fires have been responsible for massive forest fires in Uttarakhand in the past. On the other hand, careful bird monitoring can be done during ‘controlled fires’ used forcreating ‘fire lines’ against forest fire spread.

- Baiting to attract the Quails (Rieger & Walzthöny, 1993; Kalsi et al, 2007).

Comment: This will most likely be fruitless, as there are many grain/seed-eating birds and mammals in their habitat. The chances that HQ gets attracted to any bait first, are minimal. Continued baiting can also result in dependency by some animals, especially for winter migrants, who may forego foraging in favour of supplied food, which can have negative consequences in the long run. The latter report cited above, suggest the use of grain-baited photo-traps. There already exists atleast 9 camera traps (though, not grain-bait traps) in the study area of Nainital (Nainital Zoo, personal communication, September 26, 2018). This may have picked up atleast a hint of this species over the years. But, no such sightings have been made.

- Molecular/genetic analysis of egg shells/feathers from the potential habitat (Kalsi et al, 2007)

Comment: During the study, no indication of the species via the indirect means such as broken egg-shells nor shed/plucked/accidentally removed feathers were observed. All feathers in good and partially discernible state, found in the field were photographed, of which none matched illustrations or descriptions of the feathers (in Rieger & Walzthöny, 1993). Since genetic information is currently unavailable, baseline genetic information need to be extracted from museum specimens (ugently to avoid further deterioation) for comparison to field samples collected. Specialized wildlife research institutes in India and abroad could be involved in such an initiative.

#### 4.3.1. More Suggestions

Since the species has gone undetected for so long, we suggest that future explorers/scientists should focus on other potential habitats narrowed down during this study (Fig. 2 shows areas in elevational range of the species, which are understudied) and the ones suggested by Dunn et al (2015). Problem with all the surveys carried so far is that (including this one), is that it relied manual search. Buta very promising alternative has been ignored – Aerial Vehicles (UAVs/drones). Drones have been successfully used for several wildlife assessment and conservation management throughout the world, including India (with a recent establishment of a Forest Drone Force in Uttarakhand).

**Fig. 2.**
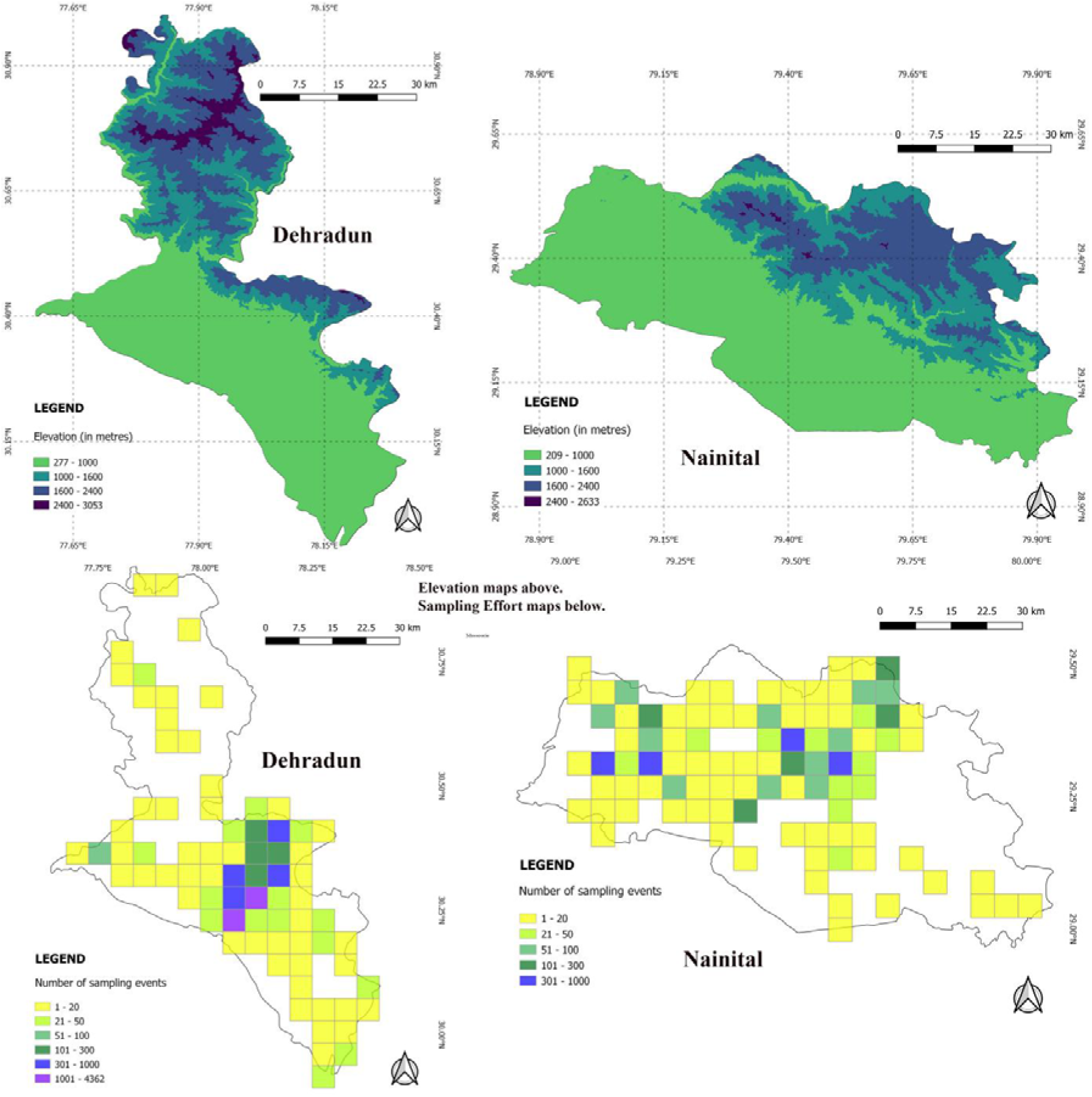
Elevation and sampling effort maps for the study area. Data for sampling efforts obtained from eBird and the current study (n = 10 541 for Dehradun and n = 4 254 for Nainital, after filtering for coordinate precision and distance falling within 5×5 Km grids).

Here is a list of benefits of using a drone instead of manual search (See Linchant et al (2015) for more details):

- These days, it is possible to procure very light-weight, silent, efficient, easy to operate and cost-effective UAVS.
- They can be remotely controlled and reach places where it was not possible for humans to search. Since the species under study is very immobile, if absolutely needed, the drones can be used to flush them, when the handler of the drone sight suspicious figures.
- Unlike humans, the drones can tolerate very cold conditions during the winter (in fact, colder conditions may give better images for the purpose).
- They can be fitted with thermal infrared-imaging, which could let handlers easily spot any signatures that bears resemblance to HQ individuals.

If UAVs is not possible at the moment, then thermal imaging hand-held cameras and/or binoculars seems like a promising option.

### 4.4. What if? – a hypothetical scenario

What if someone finally rediscovered the bird? What would be the next ideal step? Should the individual(s) reveal this marvelous news to the world? Or keep it a secret amongst a few people who are involved in the conservation of the habitat (eg: the Forest Dept. of the area)? This has been discussed by Meijaard & Nijman (2014) and Ryan & Baker (2016) by the use of decision theory and mathematical modeling. The use of the equation posited by Ryan & Baker (2016) is useful, but finding the values of the variables involved is difficult. We concur with adopting the precautionary principle (Meijaard & Nijman, 2014), atleast until we can see to it that a stable population has been established by protecting their habitat (a secrecy-based approach which has been employed before (Fitzpatrick et al, 2006)). This is due to the potential threats it faces upon public announcement.

### 4.5. Management Recommendations

Here are some recommendations for the management of the areas where the species historically occurred in/potentially occur in.

- Since the only Act for the regulation of grazing in the forest and public lands - “The Cattle Trespassers Act’ (1871) is outdated, the Task Force of Grasslands and Desert Ecosystems had called for a National Grazing Policy (Singh et al, n.d). This is very necessary as a lot of the land in the potential habitat for HQis currently under grazing pressure.
- Sterilization of domesticated/feral *C.familiaris* staying close to the potential habitats need to be carried out as they have been successful in reducing the population of free-ranging dogs (Schurer et al, 2015). This can be carried out by the help of local veterinarians and wildlife veterinarians, with the use of most ethical and efficient of Capture-Neuter-Vaccinate-Return techniques (MacFarlaine & Gibson, 2018).
- Although quite controversial, measures to reduce the growth in human population is also required (Kopnina, 2016). Awareness about family planning and proper sexual healthcare measures from local clinics can go a long way in attaining this.
- Patrol in tourist frequented areas is necessary to curb the unruly behaviour towards wildlife/in the wilderness areas. Regular cleanups will be helpful in maintaining the potential habitats.
- A cap/limit on the number of visitors allowed inside potential habitats can be set after assessing their respective carrying capacity. This also would act as a symbol showing the urgency of keeping human population under check.

## CRediT authorship contribution statement

**Paul Pop:** Data curation, Formal analysis, Investigation, Methodology, Visualization, Writing - original draft; Writing - review & editing. **Puneet Pandey:** Conceptualization, Funding acquisition, Methodology, Project administration, Supervision, Writing - review & editing. **Randeep Singh:** Conceptualization, Funding acquisition, Methodology, Project administration, Supervision, Resources, Writing - review & editing.

## Acknowledgement

This study was funded by Mohamed bin Zayed Species Conservation Fund [grant number 172517095]. We express our gratitude towards the Uttarakhand Forest Department for granting permission to work in protected areas in Nainital district. We are indebted to the people in Nainital district and Mussoorie who helped with accommodation and some of the logistics – Pratap Singh, Sajwan, Shikha, Mahesh, Deepak and Pawan being a few of them. We extend our appreciation to people who supplied us with useful data - Tim Doherty from Deakin University (supplementary material of published research) and Uttarakhand Tourism Board (tourism statistics for the state).

## Data accessibility

All bird checklists, and associated information (including coordinate data) collected during the surveys will be made available for full and free public access in eBird platform atmost by 2021.

