## Supporting Information for "Gone in 40 years, the curious case of the Himalayan Quail: an attempt at rediscovery and implications for conservation" (from another version of this preprint: version 1, 2020-07-15)

| **Table S1: Statistics of threatened Galliformes of the world** | | | | |
| --- | --- | --- | --- | --- |
| Taxa | VU | EN | CR | EX |
| Phasianidae | 24 | 11 | 3 | *1 |
| Other Galliformes | 24 | 11 | 5 | **1 |
| All Galliformes | 48 | 22 | 8 | 2 |

**Coturnix novaezelandiae* (New Zealand Quail)

***Mitu mitu* (Alagoas Curassow) in the Cracidae family (Extinct in the wild)

*Note*. Adapted from Handbook of the Birds of the World and BirdLife International digital checklist of the birds of the world. Version 4. Retrieved from http://datazone.birdlife.org/userfiles/file/Species/Taxonomy/HBW-BirdLife_Checklist_v4_Dec19.zip [.xls zipped 1 MB].

| **Table S2: Avian extinctions partly contributed by *Canis familiaris*** | |
| --- | --- |
| Scientific Name | Common Name |
| *Alopecoenas salamonis* | Thick-billed Ground-dove |
| *Coturnix novaezelandiae* | New Zealand Quail |
| *Hypotaenidia dieffenbachii* | Dieffenbach's Rail |
| *Mergus australis* | Auckland Merganser |
| *Microgoura meeki* | Choiseul Pigeon |
| *Zapornia sandwichensis* | Hawaiian Rail |

| **Table S3: Galliformes threatened by** C.familiaris | |
| --- | --- |
| Scientific Name | Common Name |
| *Aepypodius bruijnii* | Waigeo Brushturkey |
| *Eulipoa wallacei* | Moluccan Megapode |
| *Leipoa ocellata* | Malleefowl |
| *Macrocephalon maleo* | Maleo |
| *Megapodius bernsteinii* | Sula Megapode |
| *Megapodius laperouse* | Micronesian Megapode |
| *Megapodius layardi* | Vanuatu Megapode |
| *Megapodius nicobariensis* | Nicobar Megapode |
| *Megapodius pritchardii* | Tongan Megapode |
| *Cyrtonyx ocellatus* | Ocellated Quail |
| *Catreus wallichii* | Cheer Pheasant |

*Note*. Table S2 and Table S3 adapted from tables in the main text and supplementary materials, respectively, of *The global impacts of domestic dogs on threatened vertebrates*. Derived from Doherty et al (2017)^*^.

*Doherty, T. S., Dickman, C. R., Glen, A. S., Newsome, T. M., Nimmo, D. G., Ritchie, E. G., Vanak, E.T., & Wirsing, A. J. (2017). The global impacts of domestic dogs on threatened vertebrates. Biological Conservation, 210, 56-59.

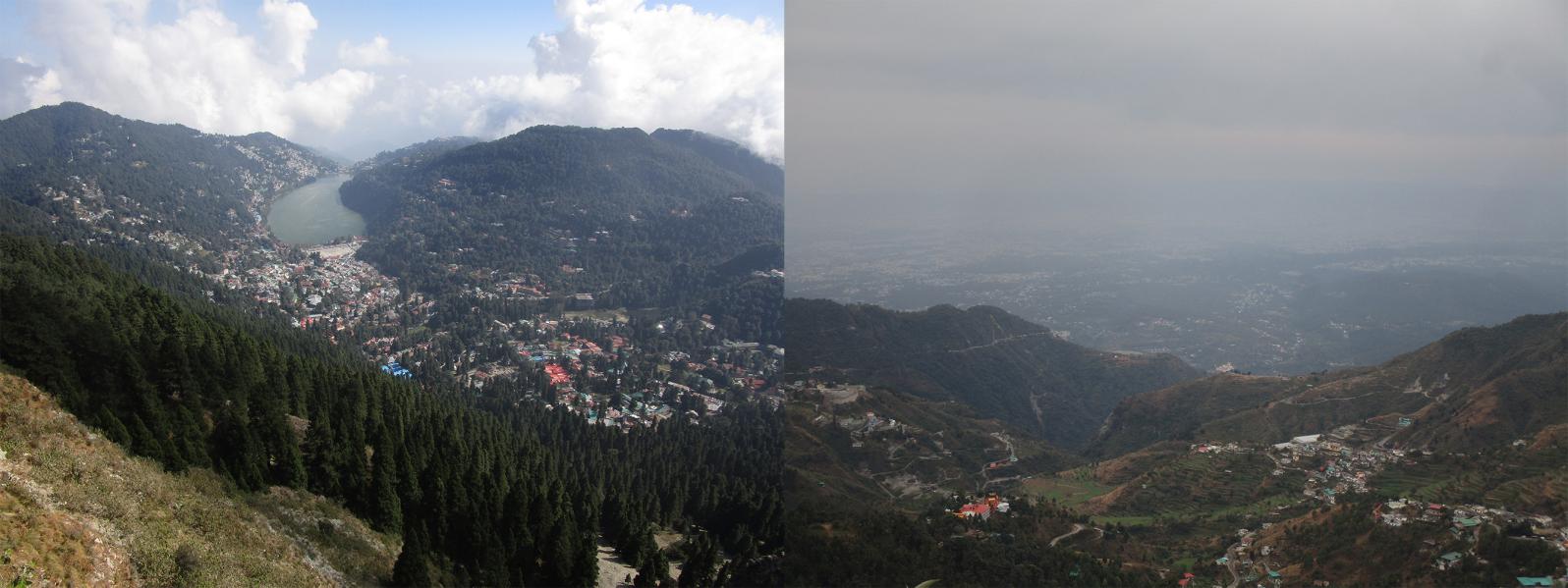

**Plate S1 – Study Area:** Nainital town in Nainital district (Left); Mussoorie town in Dehradun district (right).

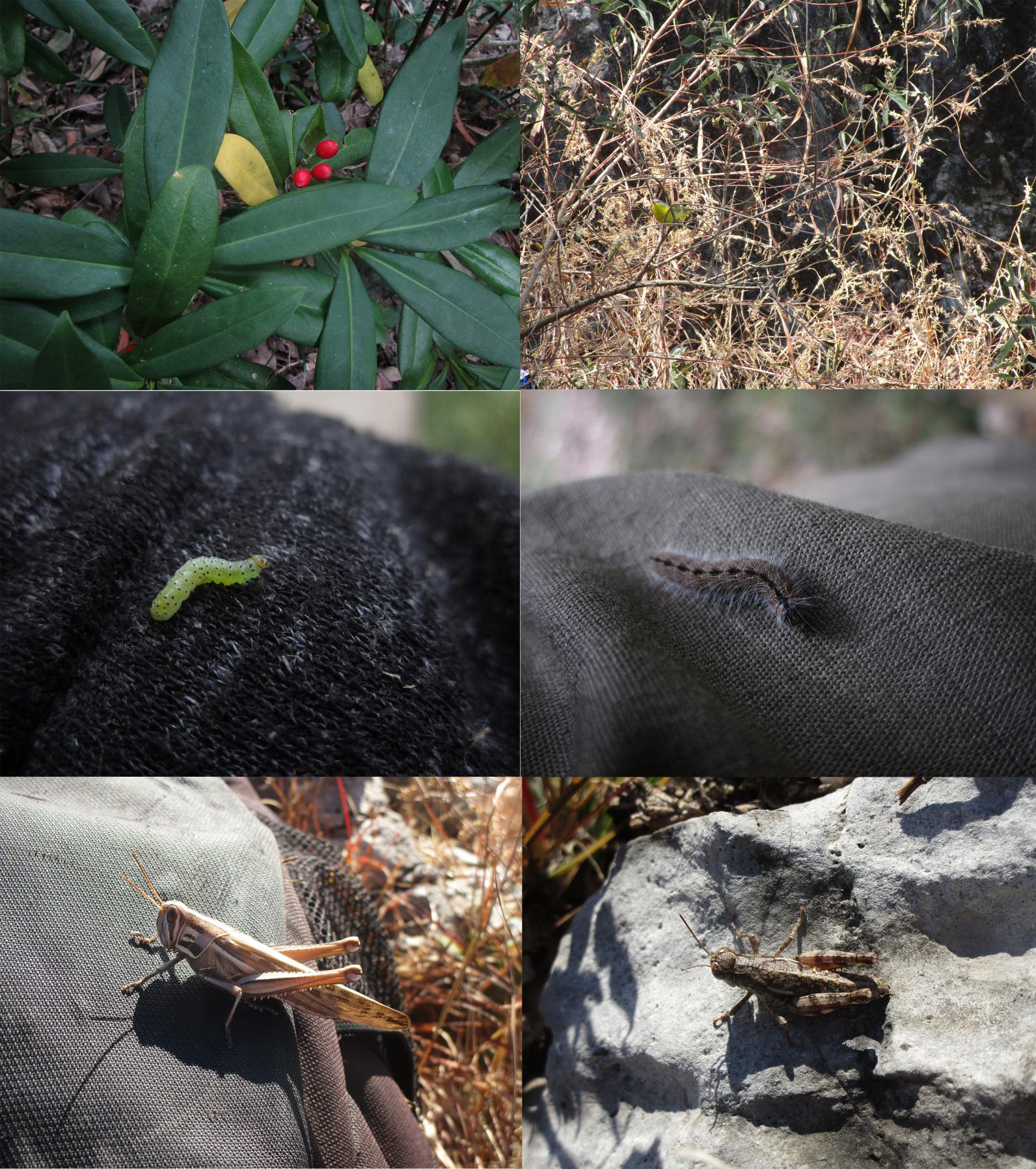

**Plate S2 – Quail Diet:** Probable components of *O.superciliosa* diet (Clockwise from top left) – a) fruits growing near the forest floor b) seed-grasses c), d), e) & f) Insects (Lepidoptera and Orthoptera).

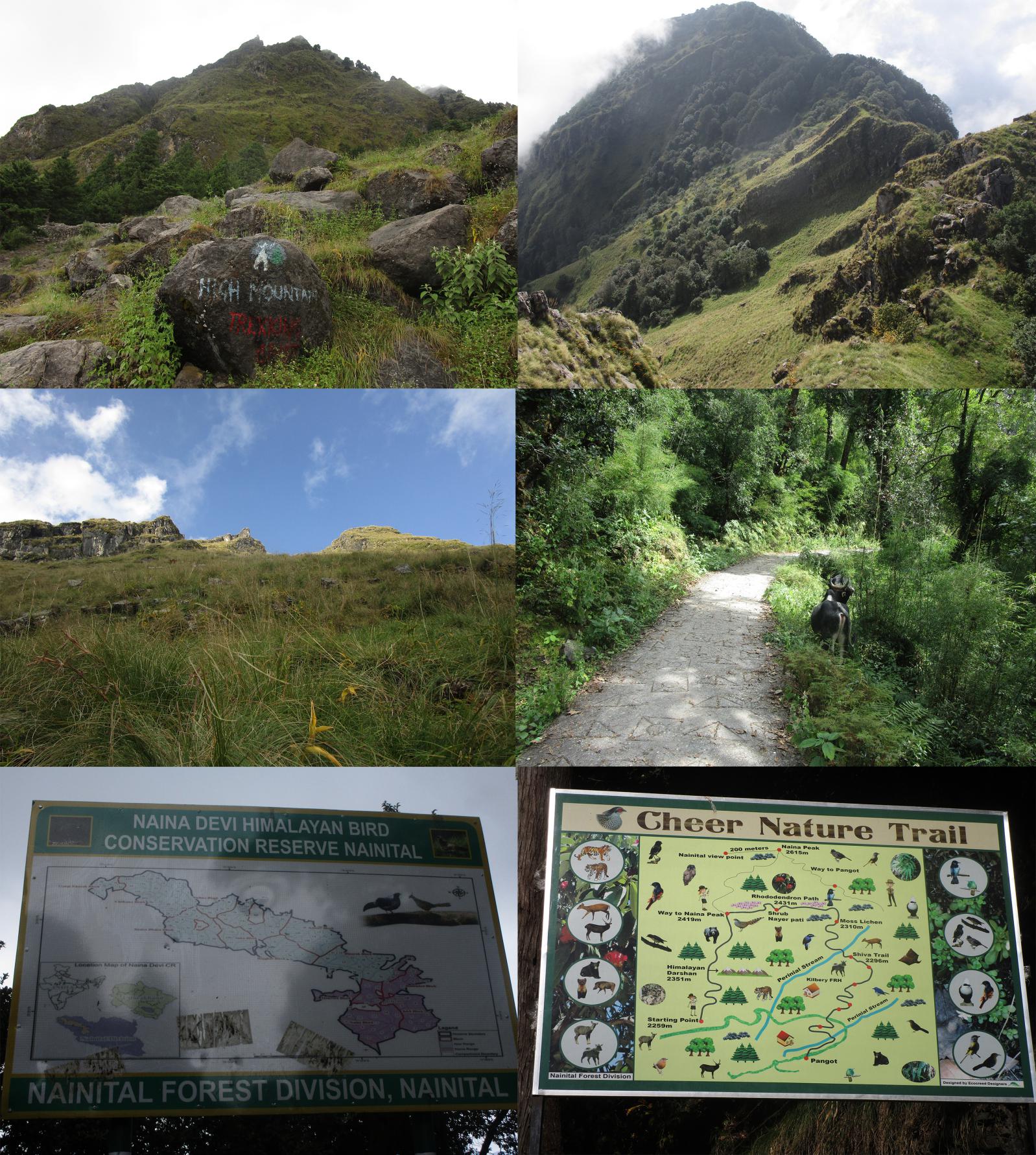

**Plate S3 – Nainital a:** (Clockwise from top left) a) Boar Peak from below; b) a view from Boar Peak; c) back route below Boar Peak; d) a pictorial map of prominent wildlife/scenic areas connected to Cheer Nature Trail; e) a map of Naina Devi Bird Conservation Reserve, close to Nainital town; f) grassy slopes at Boar Peak.

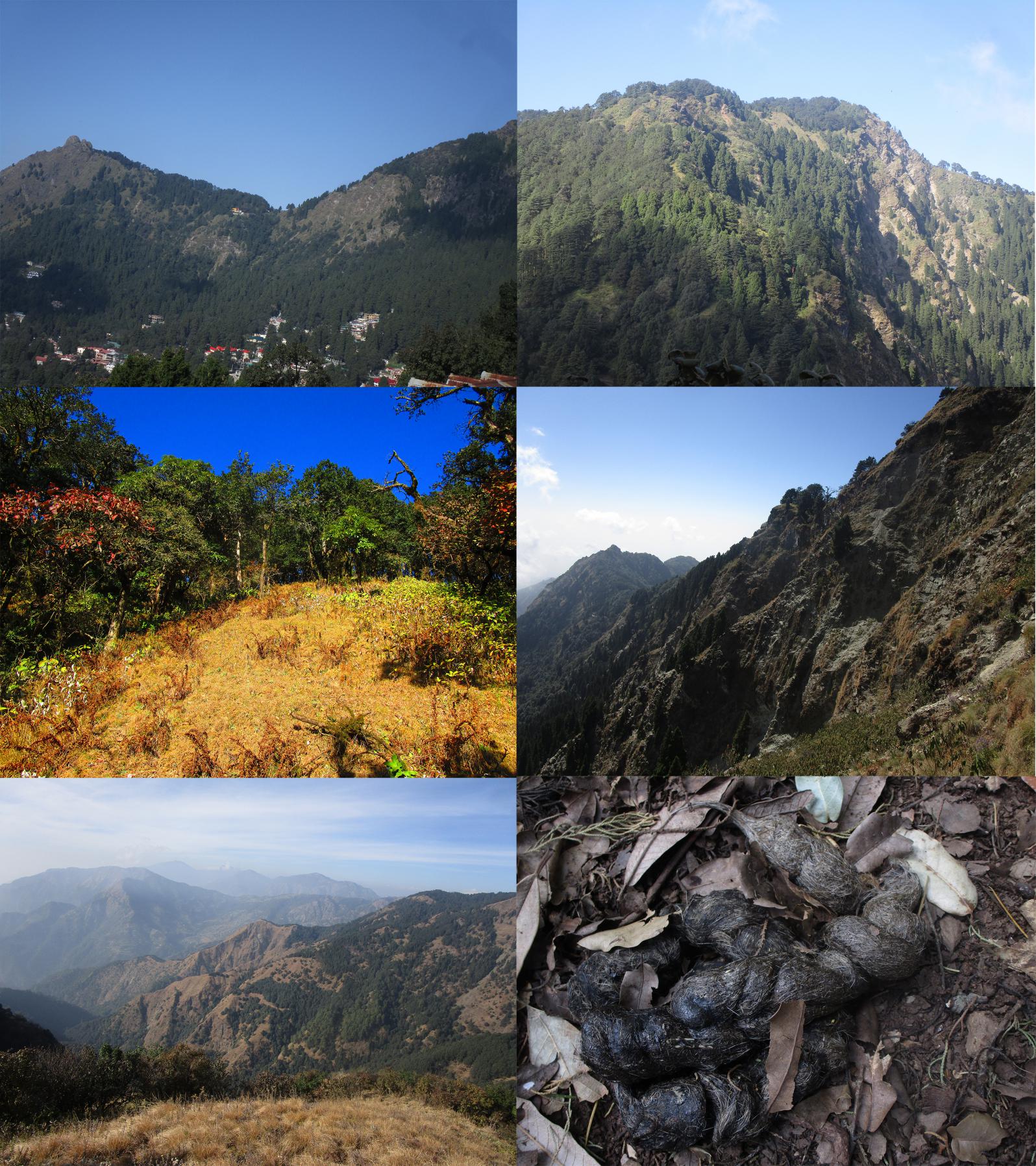

**Plate S4 – Nainital b:** (Clockwise from top left) a) Boar Peak on the top left and Naina Peak on the top right; b) Naina/China Peak; c) rocky cliff intersperse with grasses in the Naina Peak; d) Leopard scat in an isolated route near China Peak; e) distant grasslands left partially un-surveyed due to inaccessibility; f) grassy opening on a tree-filled flowery slope near Naina Peak.

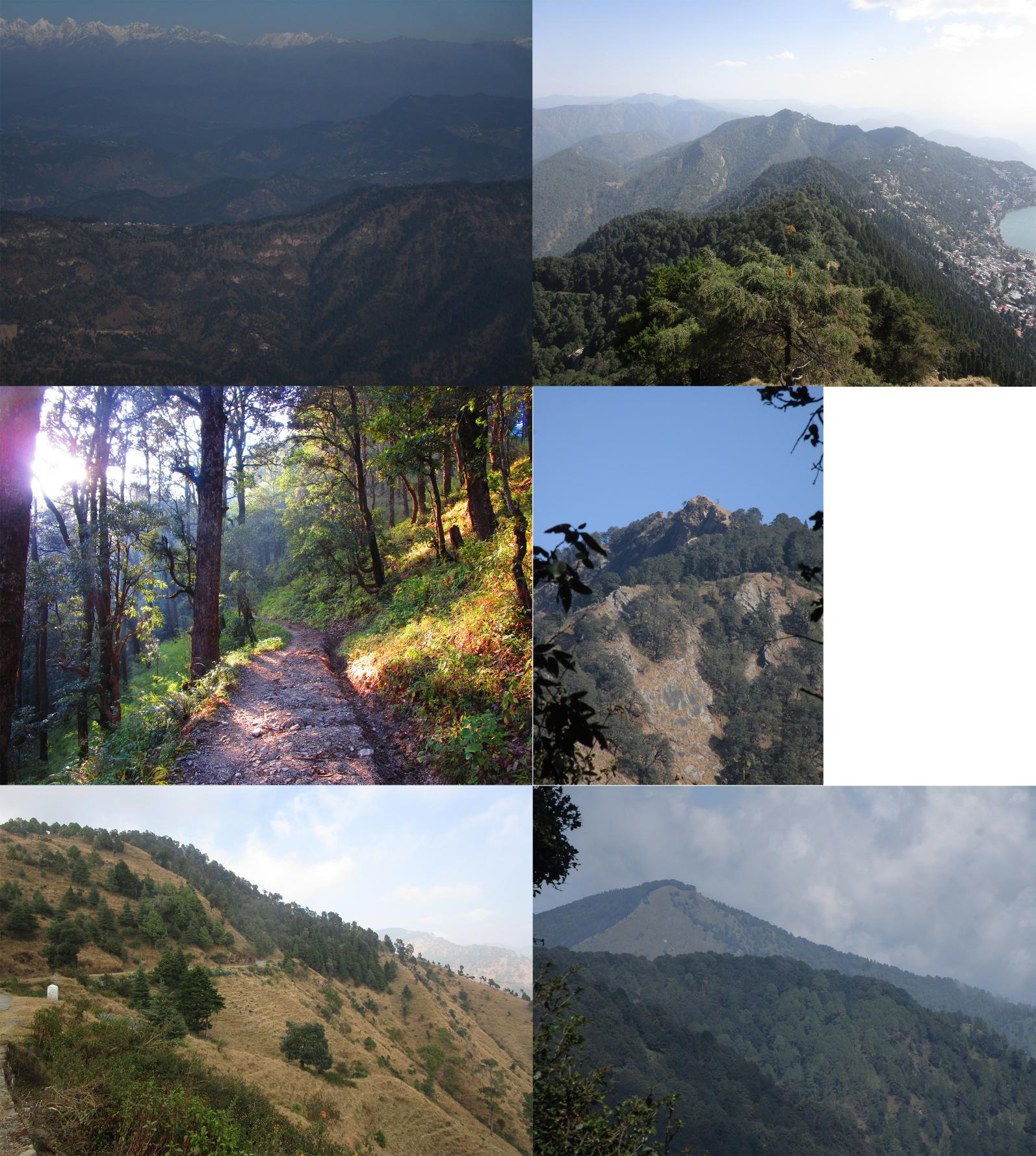

**Plate S5 – Nainital c:** (Clockwise from top left) a) Fragmented pine/oak forests and grasslands in sher-ka-danda and beyond; b) portion of Sher-ka-danda close to the town (view from China Peak); c) & d) view of grassy slopes from/inside Kahal Jungle; e) grassy slopes close to Neelkanth Hotel; f) trail route inside Kahal Jungle.

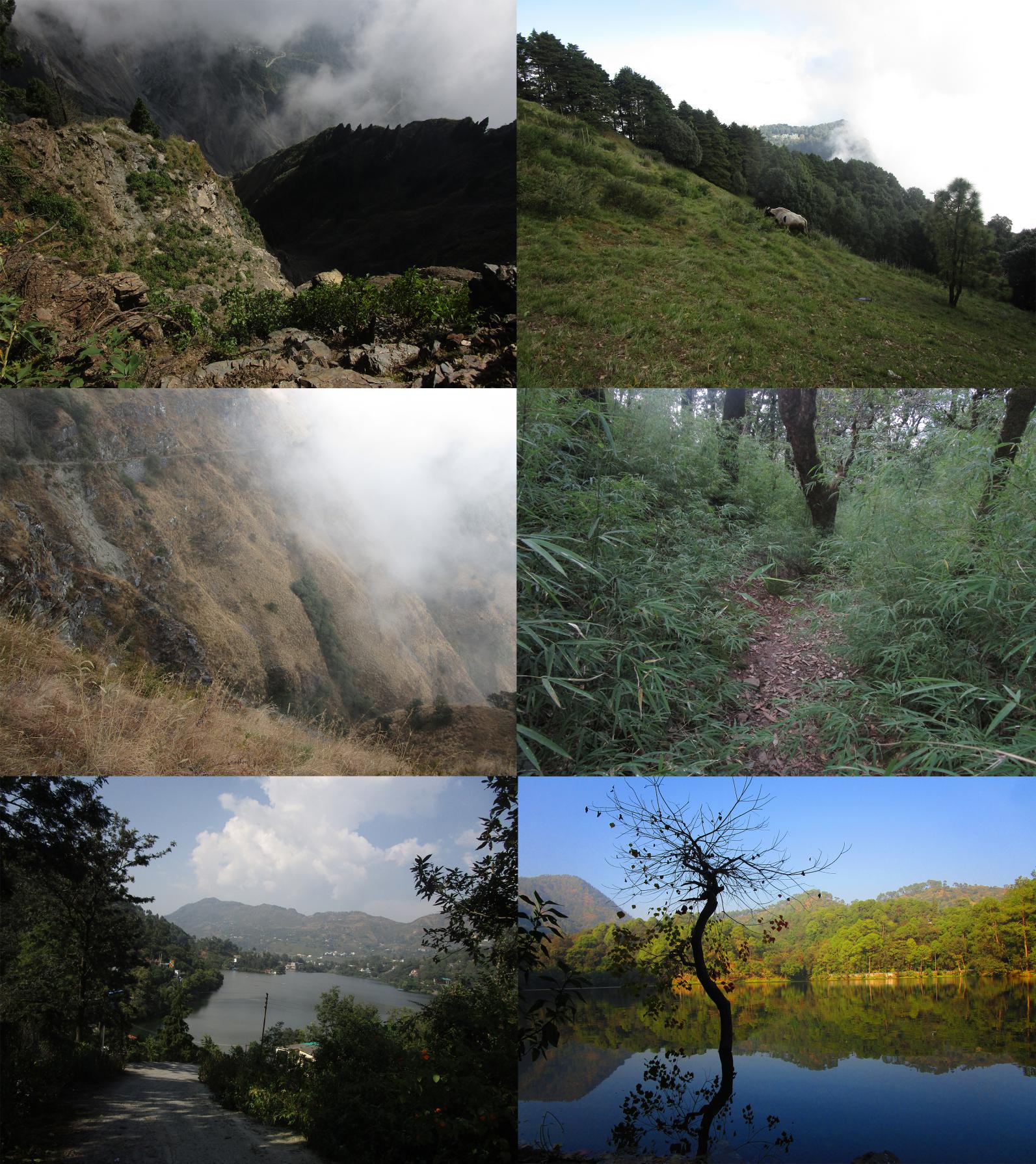

**Plate S6 – Nainital d:** (Clockwise from top left) a) Cliff-side close to Tiffin Top; b) grassland/pastureland patch near Tiffin Top; c) bamboo forest close to Tiffin Top route; d) one of seven interconnected lakes of Sattal view, with mountains in the background (highly disturbed area); e) Naukuchiyatal with mountains in the background (highly disturbed area); f) Cheer Point.

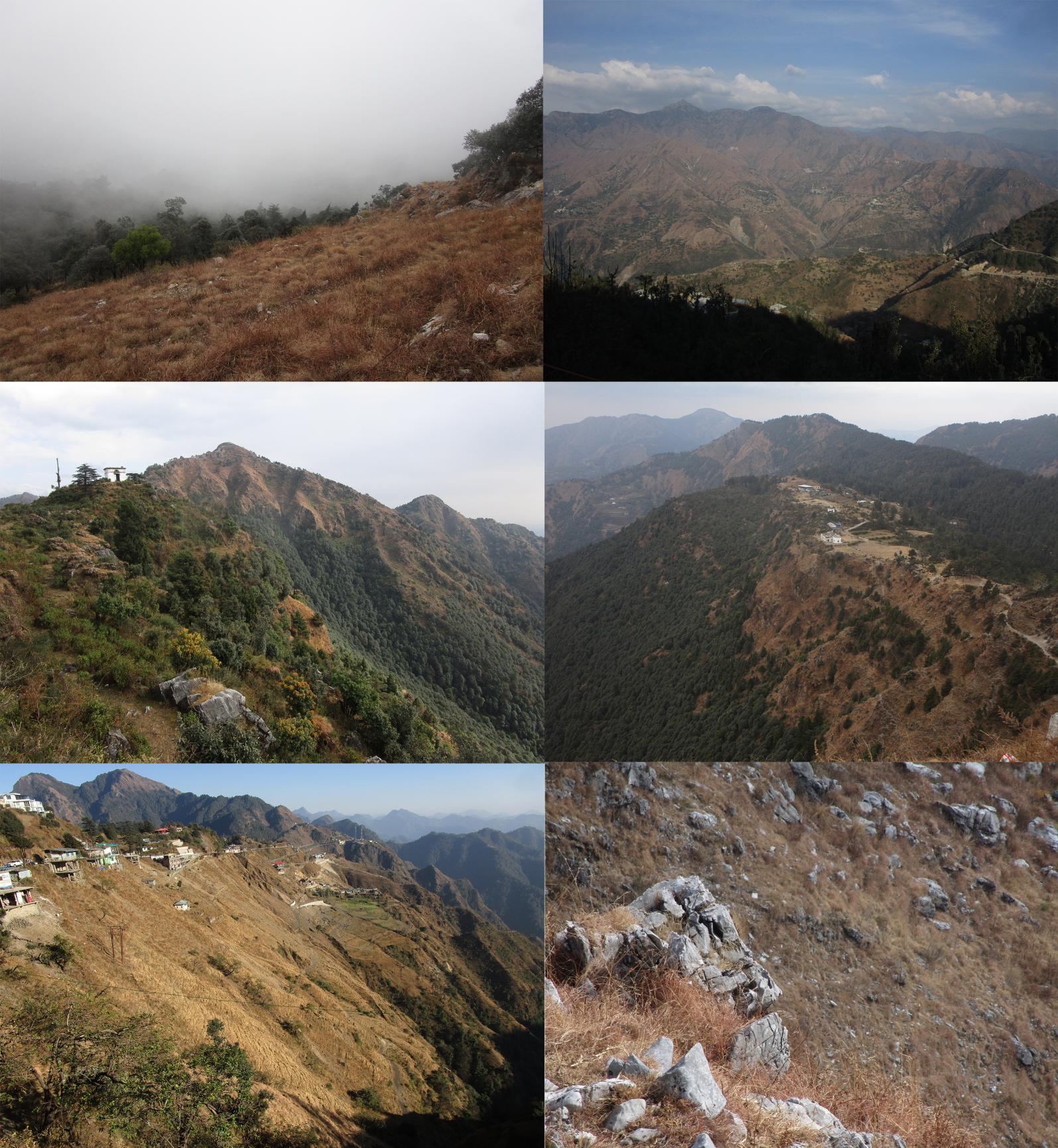

**Plate S7 – Dehradun a:** (Clockwise from top left) a) Grassy patch inside Jabarkhet Nature Reserve; b) A view of the valleys and mountains in Dehradun from Mussoorie-Kempty road; c) view from slope of George Everest; d) *Prunella himalyana* (Altai Accentor) camouflaged against the rock/grass background at George Everest; e) Grassland/pasturelands in Masrana village; f) George Everest peak (top middle).

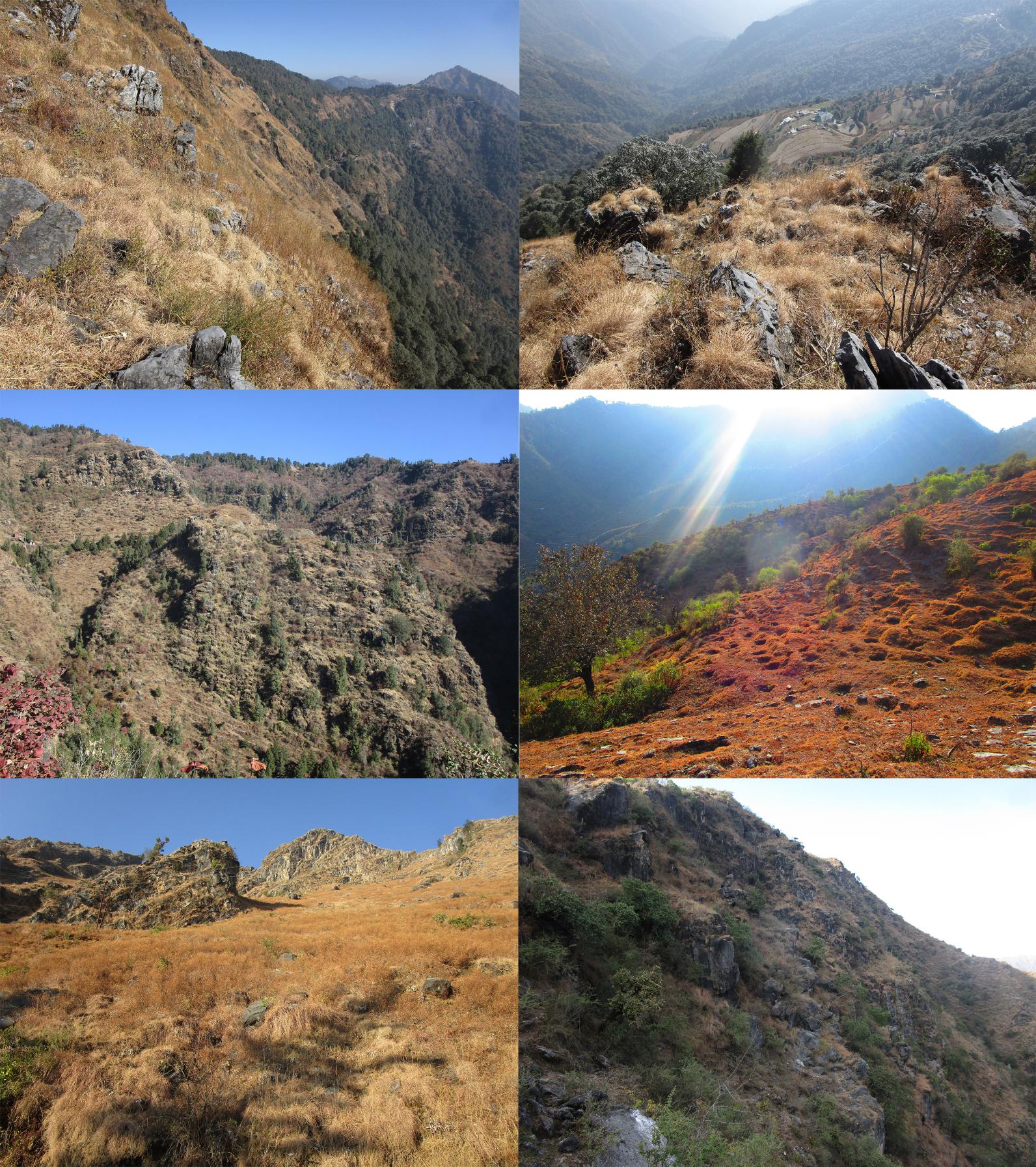

**Plate S8 – Dehradun b:** (Clockwise from top left) a) View of Cloud Ends view point; b) farms in the vicinity of Benog Wildlife Sanctuary; c) Grassy slopes of the peak adjacent to Kempty falls; d) Cliffside of the same peak; e) & f) Relatively undisturbed long-grass habitats between Mussoorie and Dhanolti.

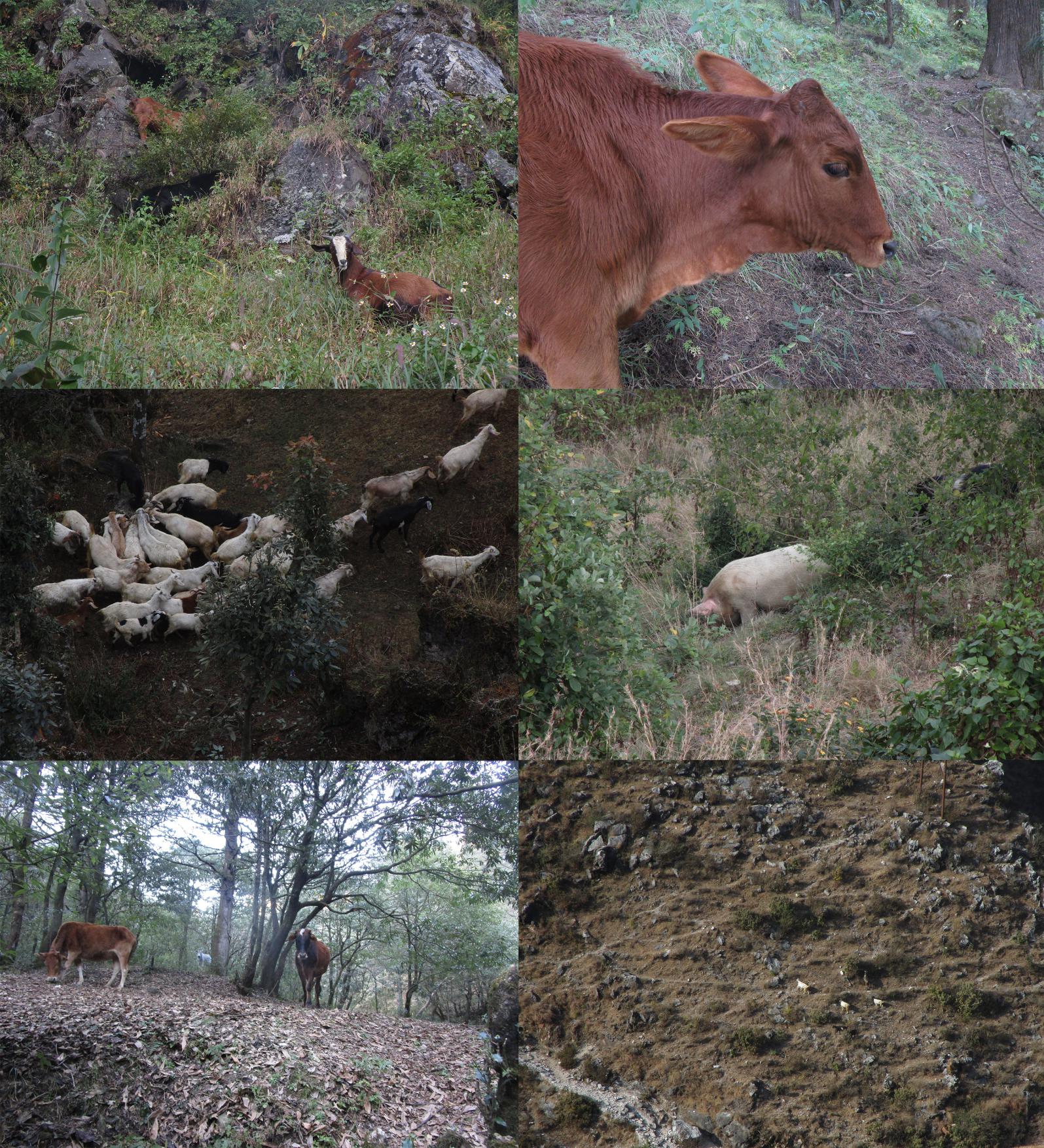

**Plate S9 – Grazing and Foraging:** (Clockwise from top left) a) Goats browsing at Ayarpatta side, in Nainital; b) a cow foraging on the slopes of Boar Peak, Nainital; c) a sounder of domestic pigs foraging in the grassy slopes near Neelkanth Hotel, Nainital; d) a herd of goat foraging in the slopes of the peak adjacent to Kempty Falls e) a cow-herd grazing in the buffer zone of Benog Wildlife Sanctaury; f) a large flock of goats browsing at the slopes of George Everest.

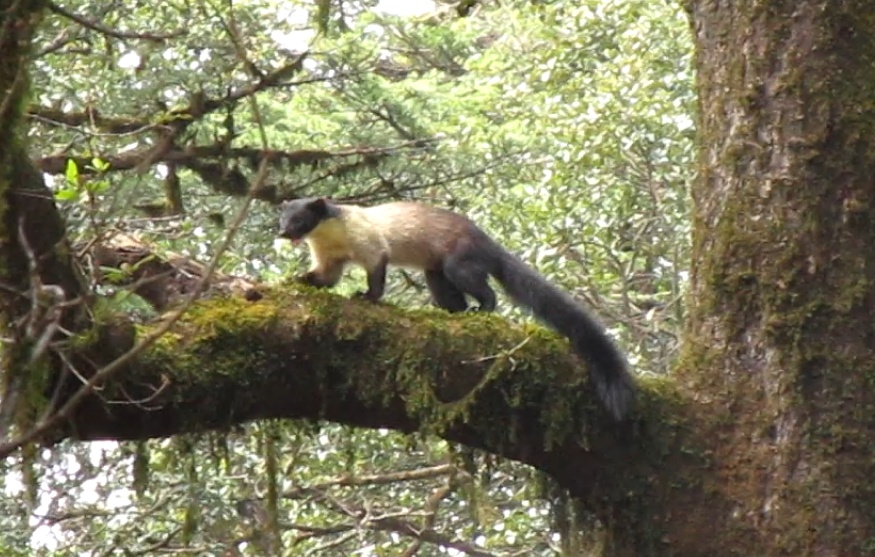

**Plate S10 - A Predator:** *M.flavigula* on a tree-top inside an oak forest near China Peak, part of a pack of three.

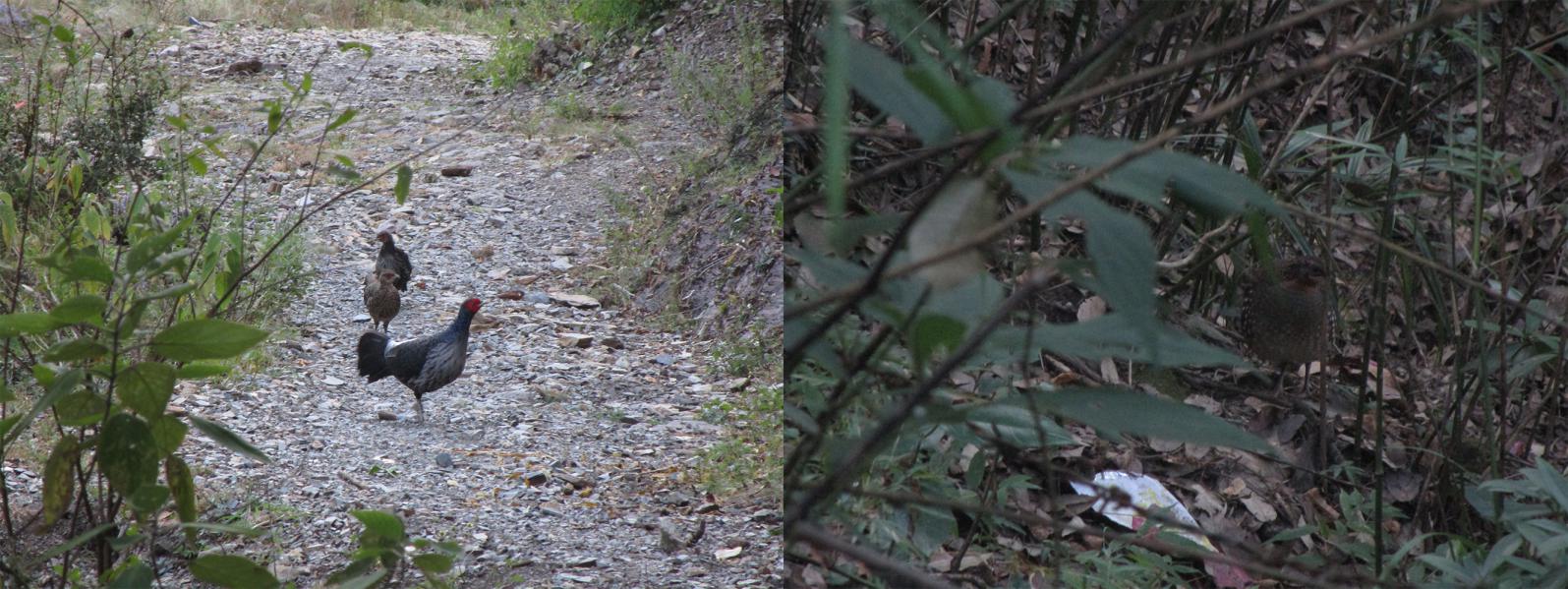

**Plate S11 - The Hunted:** Part of a covey of *L.leucomelanos* (male and female) (left); *A.torqueola* (female) (right)

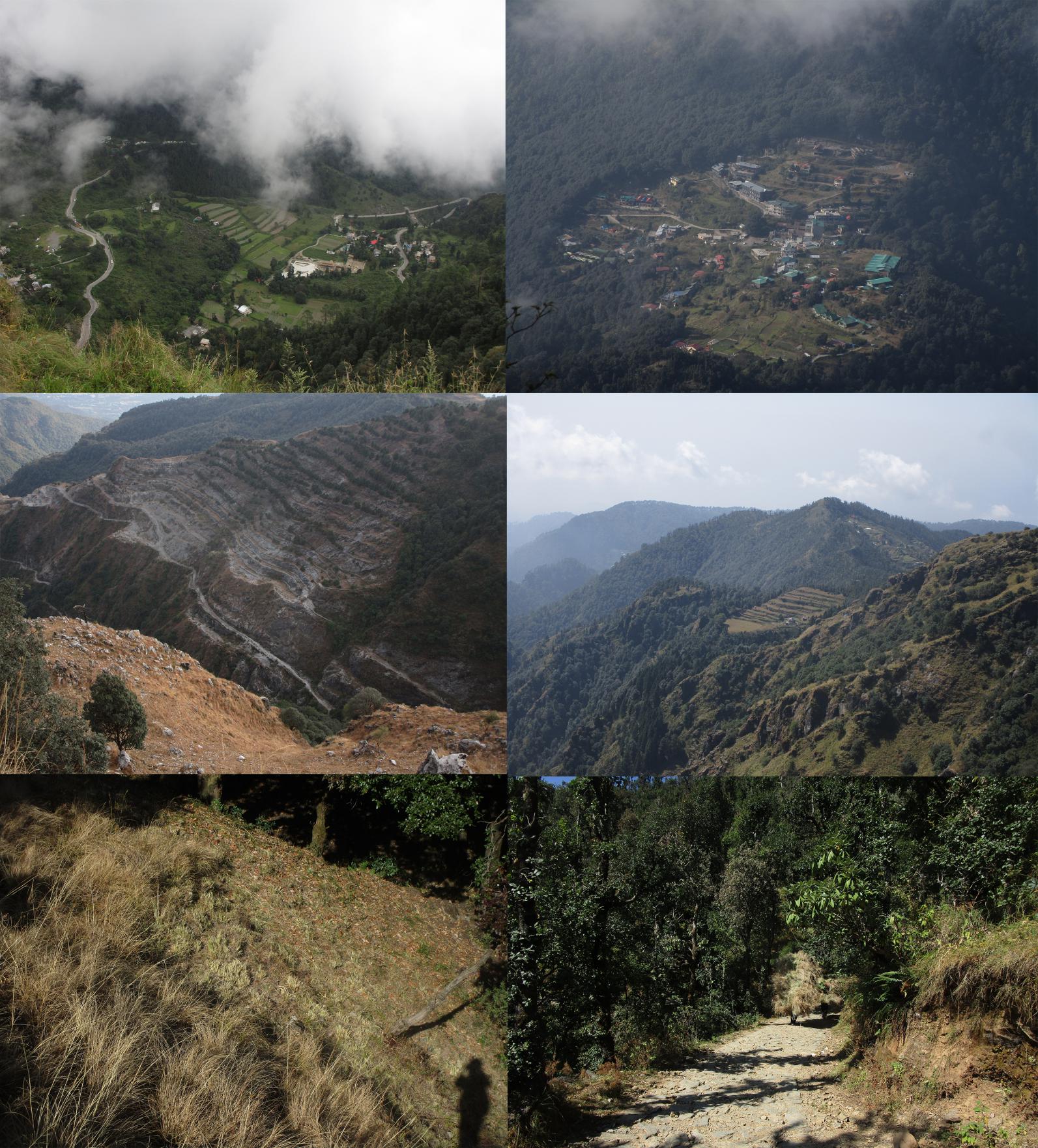

**Plate S12 – Landuse Changes a:** (Clockwise from top left) a) Agricultural lands in the valley below Nainital town; b) settlements in clear-felled lands close to Khurpa tal; c) agricultural field close to the top, near Boar Peak; d) grass collection near Boar Peak; e) difference between uncut and cut grass, side-by-side at Boar Peak; f) Mined slope opposite to George Everest.

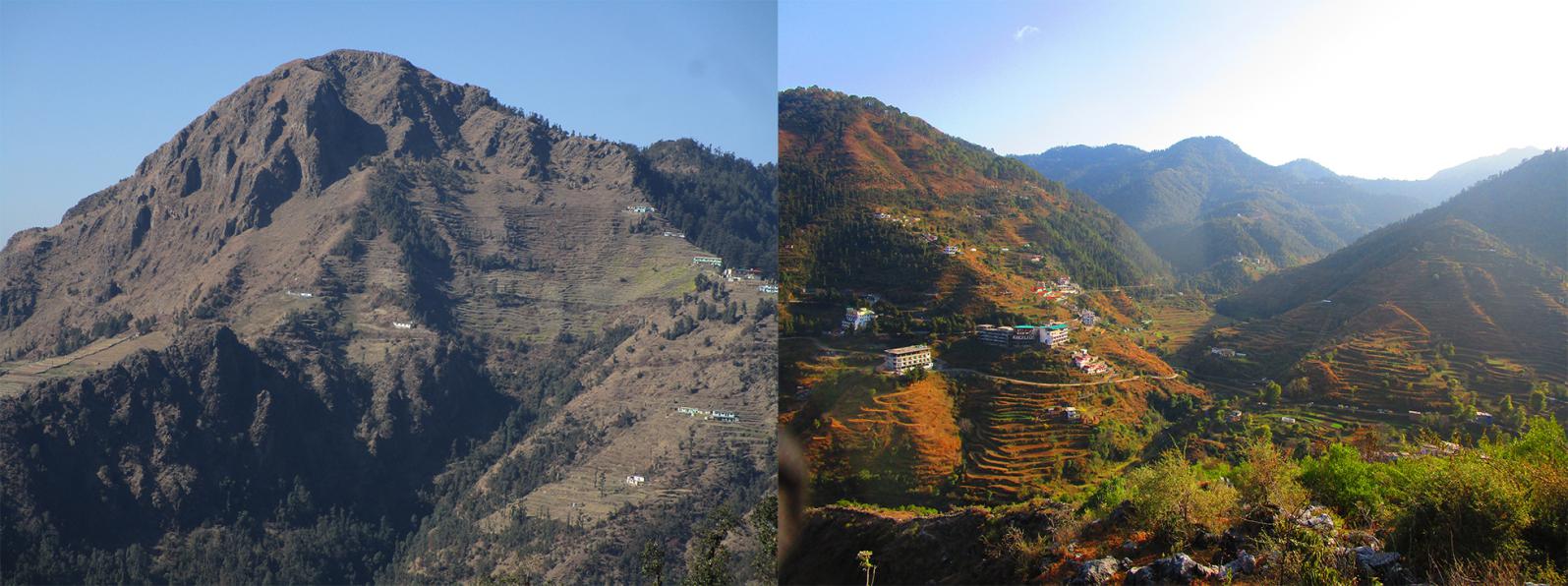

**Plate S13 – Landuse Changes b:** a) terrace farming on the mountain-slopes of Dhanolti (left); b) step cultivation in Ramnagar, Kempty region (right).

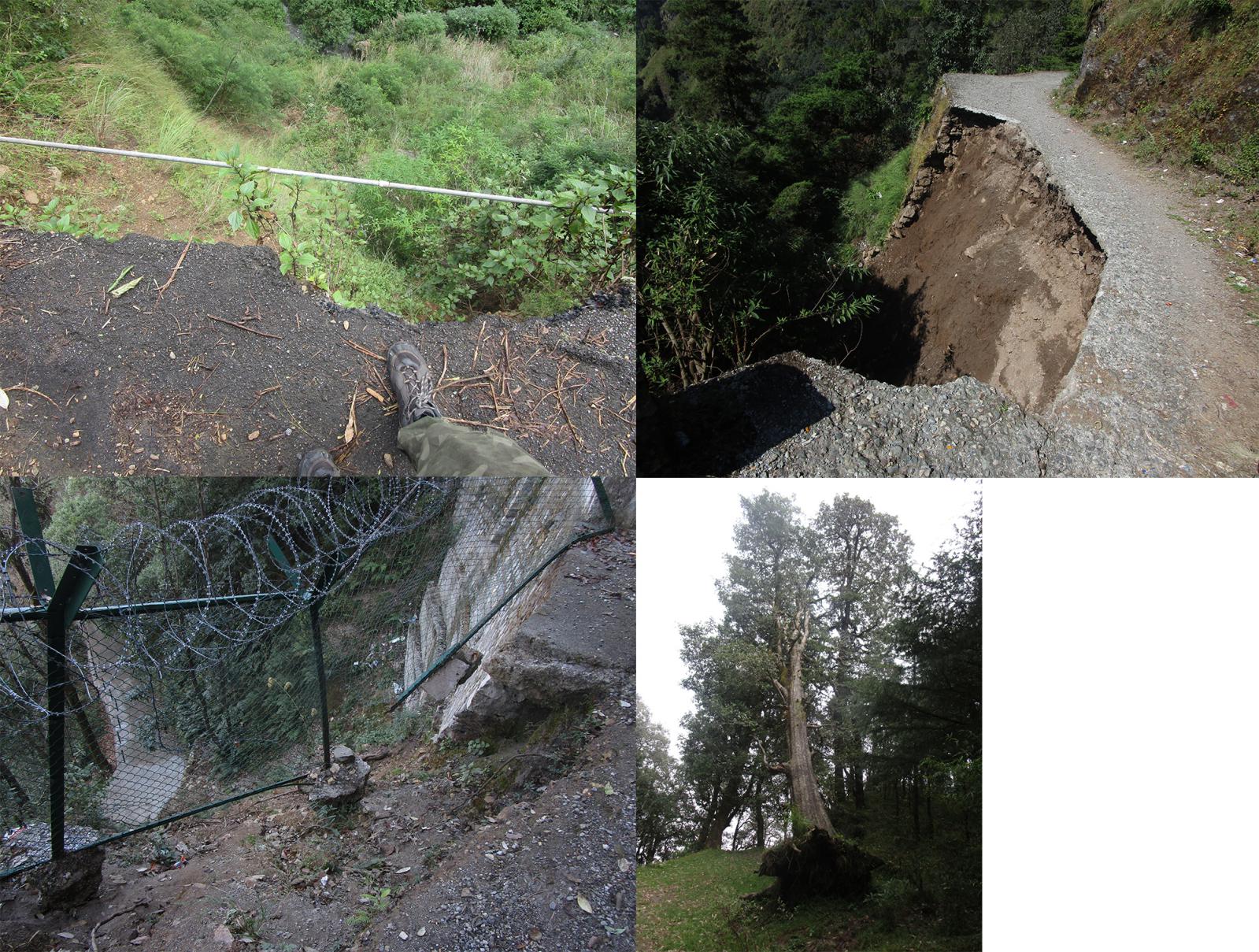

**Plate S14 – Biogeochemical Hazards:** Continuous shifting of soil and matter due to thawing, erosion, wind, rain and other processes causes structural damage to human-made structures and flora. (Clockwise from top left) a) damaged concrete roadside in Sattal; b) damaged footpath in the backside of Boar peak; c) uprooted tree near Tiffin Top; d) damaged fence close to Hathipaon, Mussoorie.

oc

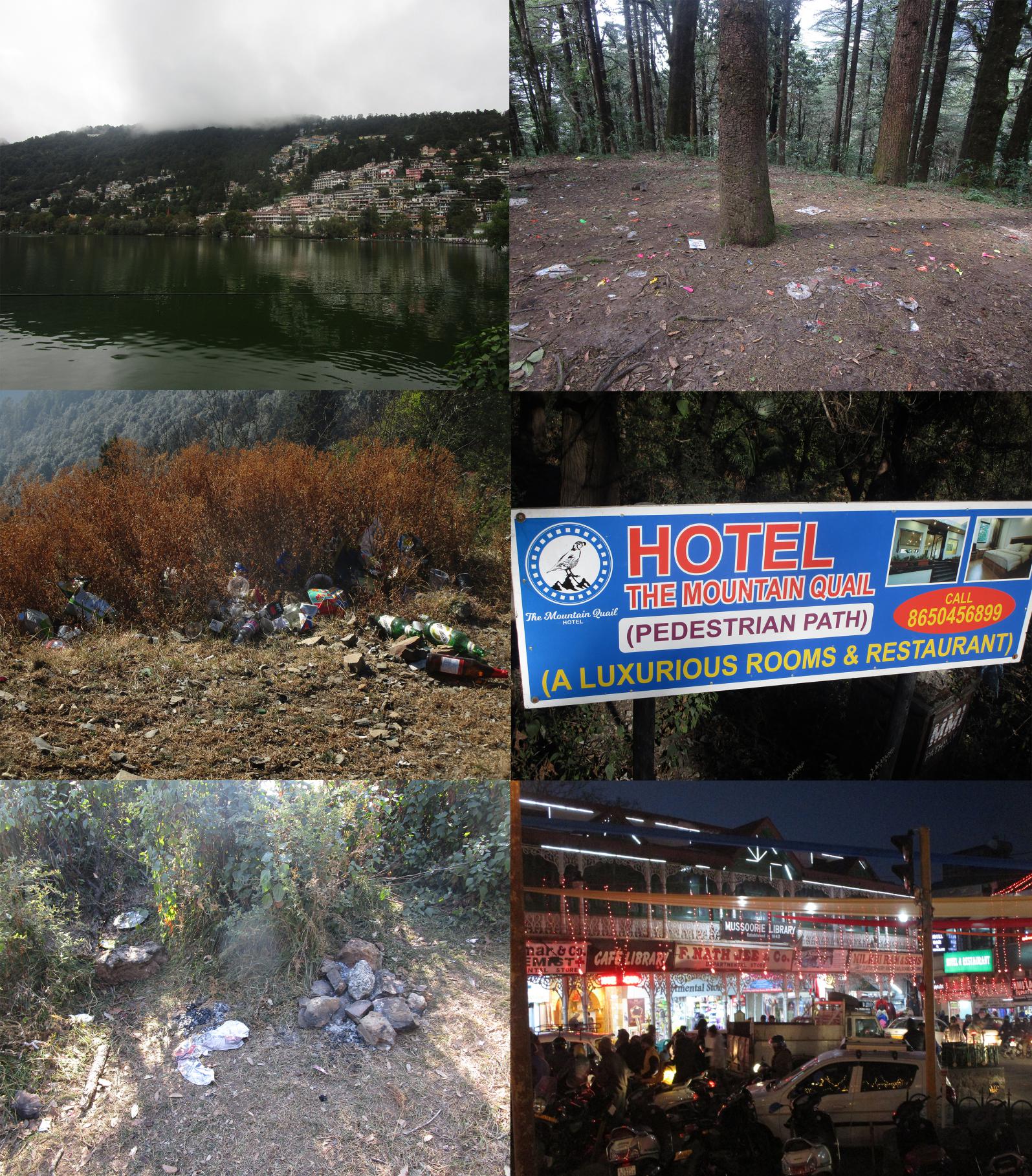

**Plate S15 – Effects of Tourism a:** (Clockwise from top left) a) Resorts in Nainital town - there has been a tremendous growth in number of resorts in the last century and it’s still growing; b) littering after partying inside the pine forest on the way to Boar Peak c) while trying to cash on what’s Mussoorie is known for, the hotel management didn’t even to bother to search for the right picture of *O.superciliosa* aka Mountain Quail, but instead used an illustration of *Callipepla californica* (California Quail) as their logo – a Least Concern (IUCN) species found only in western North America d) heavy influx of tourists in Mussoorie in the end of December; e) fire started at the Cloud Ends point- potential cause of fires, especially in the summers; f) alcohol bottles, and other trash strewn across a slope in the Chamba-Mussoorie road.

oc

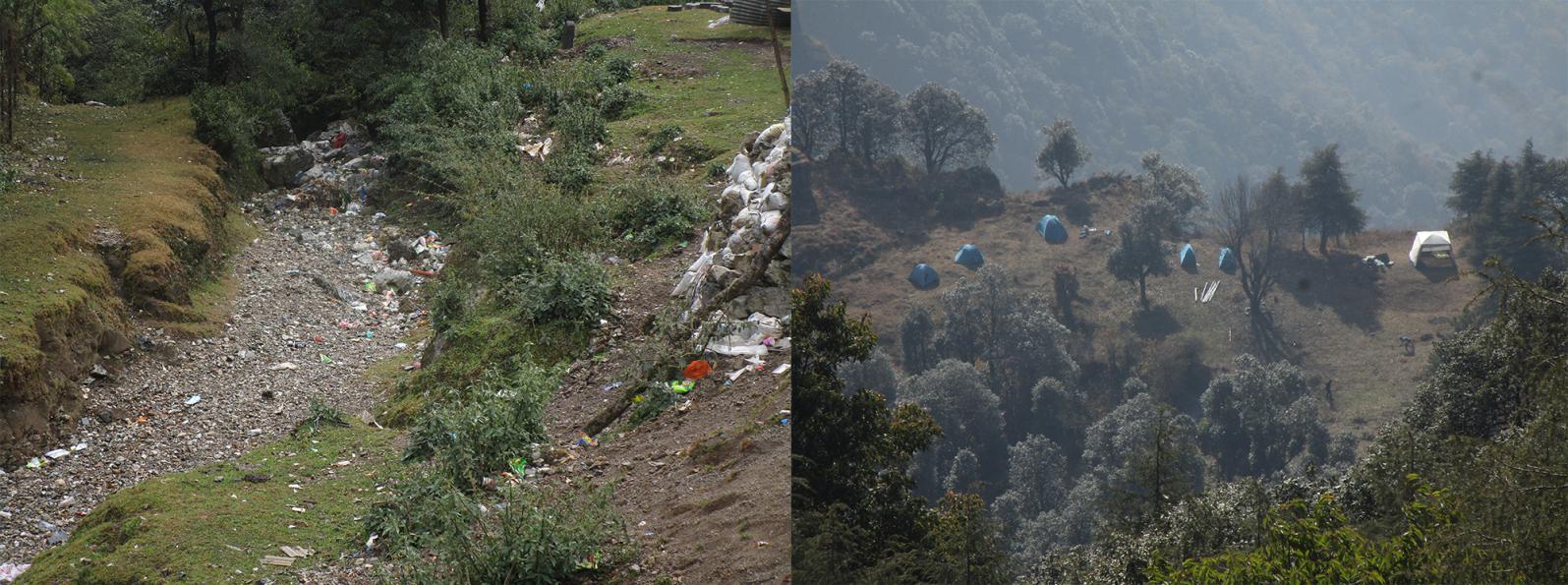

**Plate S16 – Effects of Tourism b:** a) plastic discards in the stream bed in the base of George Everest (left); b) camping in clear-felled lands close to George Everest (right).

**Figure S1. Population growth of three different towns/cities in the potential habitat of *O.superciliosa* from date of establishment to present**.

Population

| Year | Nainital | Mussoorie |
| --- | --- | --- |
| 1827 |  | 100 |
| 1839 | 100 | 1031 |
| 1901 | 6903 | 6461 |
| 1911 | 10270 | 8052 |
| 1921 | 11230 | 9702 |
| 1931 | 10673 | 6116 |
| 1941 | 10118 | 7172 |
| 1951 | 13093 | 8550 |
| 1961 | 16080 | 11238 |
| 1971 | 25167 | 20389 |
| 1981 | 26093 | 18233 |
| 1991 | 30951 | 29629 |
| 2001 | 39911 | 29329 |
| 2011 | 42775 | 33657 |
| 2021 | 52425 | 40486 |
| 2031 | 64252 | 48701 |

*Note*. Data for 20^th^ and 21^st^ century from ORGI (2013)^[1]^. Initial population taken as 100 (arbitrary reference), and modeled accordingly (1901-2011 are census data, and values for 2021 and 2031 are projections based on average decadal growth rates). Nainital and Mussoorie can be said to have been founded/established as towns in 1839 by Mr P. Barron and in 1827 by Captain Young, respectively (Rieger & Walzthöny, 1993). The population before this time was composed mostly of few farmers (let’s assume ≤100) during the founding years. Office of the Registrar General & Census Commissioner, India reports that the population for Nainital city was 42,775 (data for Nagar Palika Parishad/Municipal Board and Nainital Cantonment Board have been pooled (which was built after the census of 1921), since they are both located close by, and the population of the latter is significant enough) in 2011 (2011a, 2011b)^[2][3]^ and that of Mussoorie (Nagar Palika Parishad/Municipal Board + Landaur Cantonment Board, for the same reasons as above) was 33,657 (ORGI, 2011c, 2011 d)^[4][5]^.

^[1]^ Office of the Registrar General & Census Commissioner (2013). *Town Directory: Uttaranchal: Table for Uttarakhand* [Data file]. Retrieved from <http://censusindia.gov.in/DigitalLibrary/Tables.aspx>

^[2]^ Office of the Registrar General & Census Commissioner (2011a). *Nainital Population Census 2011-2019*. Retrieved from <https://www.census2011.co.in/data/town/800331-nainital-uttarakhand.html>

^[3]^ Office of the Registrar General & Census Commissioner (2011b). *Nainital Population Census 2011-2019*. Retrieved from <https://www.census2011.co.in/data/town/800332-nainital-uttarakhand.html>

^[4]^ Office of the Registrar General & Census Commissioner (2011c). *Mussoorie Population Census 2011 – 2019*. Retrieved from <https://www.census2011.co.in/data/town/800307-mussoorie-uttarakhand.html>

^[5]^ Office of the Registrar General & Census Commissioner (2011d). *Landaur Population Census 2011 – 2019*. Retrieved from <https://www.census2011.co.in/data/town/800308-landaur-uttarakhand.html>

**Figure S2. Growth of tourists/visitors in two different towns/cities in the potential habitat of *O.superciliosa* for the past decade**.

No. of visitors (in lakhs/100 000)

| Year | Nainital | Mussoorie |
| --- | --- | --- |
| 2009 | 0.89079 | 11.0257 |
| 2010 | 0.83609 | 11.048 |
| 2011 | 8.43815 | 11.2029 |
| 2012 | 9.06331 | 12.0529 |
| 2013 | 7.44218 | 10.7598 |
| 2014 | 7.58123 | 13.967 |
| 2015 | 8.15805 | 27.6927 |
| 2016 | 8.73395 | 27.9512 |
| 2017 | 9.18652 | 27.9597 |
| 2018 | 9.33657 | 28.7203 |
| 2019 | 9.41126 | 29.8691 |
| 2020 | 9.48655 | 31.0638 |
| 2021 | 9.56244 | 32.3064 |

Data from 2009-2018 from Uttarakhand Tourism Development Board, Government of India (2019). In Mussoorie, the number of visitors stood at 11.02 lakh in 2009, which ascended to a staggering 28.72 lakh by 2018. Data for 2019-2021 modelled using average annual growth rate (excluding the outlier caused by a large jump from 2010 to ‘11 in Nainital and an apparent jump from 2014 to ‘15 in Mussoorie). There may be some discrepancies in the data, due to difference in data collection methodologies or other factors. So, caution is taken to adjust for it, in the projection. The estimated number of tourists for Nainital in 2019 was 9.41 lakh, whereas the estimate for Mussoorie was 29.86 lakh or 29.07 lakh (if only the growth rates from 2015 to ’18 are considered). The projections for 2020 and 2021 in both locations is most likely not going to be reached due to the Covid-19 pandemic. In fact, in these two years, there is likely to be a steep decline due to the lockdown which will be beneficial to the habitats there. But when the lockdown is completely lifted, there is likely to be a sudden surge of visitors (as seen in China).

| **Table S4: Checklist of birds seen in the study area with migratory and conservation statuses** | | | | | | |
| --- | --- | --- | --- | --- | --- | --- |
| **Scientific Name** | **Common Name** | **Sighted at** | | **Migratory Status** | **IUCN status** | **IWPA status** |
|  |  | **Nainital District** | **Mussoorie District** |  |  |  |
| **Galliformes (5)** | | | | | | |
| **Phasianidae (5)** | | | | | | |
| *Arborophila torqueola* (Valenciennes, 1825) | Common Hill Partridge | × | × | R | LC | Schedule IV |
| *Catreus wallichii* (Hardwicke, 1827) | Cheer Pheasant | × |  | R | VU | Schedule I |
| *Coturnix coturnix* (Linnaeus, 1758) | Common Quail | × |  | WV | LC | Schedule IV |
| *Francolinus francolinus* (Linnaeus, 1766) | Black Francolin | × |  | R | LC | Schedule IV |
| *Lophura leucomelanos* (Latham, 1790) | Kalij Pheasant | × | × | R/AM | LC | Schedule IV |
| **Columbiformes (2)** | | | | | | |
| **Columbidae (2)** | | | | | | |
| *Columba livia* (J.F. Gmelin, 1789) | Blue Rock Pigeon | × | × | R | LC | Schedule IV |
| *Streptopelia turtur* (Linnaeus, 1758) | Oriental Turtle Dove | × |  | R | LC | Schedule IV |
| **Caprimulgiformes (3)** | | | | | | |
| **Caprimulgidae (1)** | | | | | | |
| *Caprimulgus jotaka*Temminck & Schlegel, 1844 | Grey Nightjar | × |  | SV | LC | Schedule IV |
| **Apodidae (2)** | | | | | | |
| *Aerodramus brevirostris* (Horsfield, 1840) | Himalayan Swiftlet | × |  | R | LC | Schedule IV |
| *Appus affinis* (J.E. Gray, 1830) | House Swift | × |  | R | LC | Schedule IV |
| **Cuculiformes (1)** | | | | | | |
| **Cuculidae (1)** | | | | | | |
| Cuculus poliocephalus Latham, 1790 | Lesser Cuckoo | × |  | SV | LC | Schedule IV |
| **Pelecaniformes (1)** | | | | | | |
| **Phalacrocoracidae (1)** | | | | | | |
| *Phalacrocorax carbo* (Linnaeus, 1758) | Great Cormorant | × |  | WV | LC | Schedule IV |
| **Accipitriformes (7)** | | | | | | |
| **Accipitridae (7)** | | | | | | |
| *Accipiter nisus* (Linnaeus, 1758 ) | Eurasian Sparrowhawk | × |  | R/WV | LC | Schedule I |
| OR Accipiter virgatus (Temminck, 1822) | OR Besra |  |  | R | LC | Schedule I |
| *Aquila nipalensis* (Hodgson, 1833) | Steppe Eagle | × | × | WV | EN | Schedule I |
| *Gypaetus barbatus* (Linnaeus, 1758) | Bearded Vulture |  | × | R | NT | Schedule I |
| *Gyps fulvus* (Hablizl, 1783) | Griffon Vulture | × | × | WV | LC | Schedule IV |
| *Gyps himalayensis* Hume, 1869 | Himalayan Vulture | × | × | R | NT | Schedule IV |
| *Milvus migrans* (Boddaert, 1783) | Black Kite | × | × | R | LC | Schedule I |
| *Nisaetus nipalensis* Hodgson, 1836 | Mountain Hawk Eagle |  | × | R | LC | Schedule I |
| **Strigiformes (1)** | | | | | | |
| **Strigidae (1)** | | | | | | |
| *Glaucidium brodiei* (E. Burton, 1836) | Collared Owlet | × |  | R | LC | Schedule IV |
| **Piciformes (10)** | | | | | | |
| **Picidae (8)** | | | | | | |
| *Chrysophlegma flavinucha* (Gould, 1832) | Greater Yellow-naped Woodpecker | × |  | R | LC | Schedule IV |
| *Dendrocopos auriceps* (Vigors, 1831) | Brown-fronted Woodpecker | × | × | R/AM | LC | Schedule IV |
| *Dendrocopos himalayensis*(Jardine & Selby, 1831) | Himalayan Woodpecker | × | × | R | LC | Schedule IV |
| *Dendrocopos hyperythrus* (Vigors, 1831) | Rufous-bellied Woodpecker | × | × | R | LC | Schedule IV |
| *Picumnus innominatus* E. Burton, 1836 | Speckled Piculet |  | × | R | LC | Schedule IV |
| *Picus canus* J.F. Gmelin, 1788 | Grey-headed Woodpecker | × | × | R | LC | Schedule IV |
| *Picus chlorolophus* Vieillot, 1818 | Lesser Yellow-naped Woodpecker | × | × | R | LC | Schedule IV |
| *Picus squamatus* Vigors, 1831 | Scaly-bellied Woodpecker | × | × | R | LC | Schedule IV |
| **Ramphastidae (2)** | | | | | | |
| *Psilopogon asiaticus* (Latham, 1790) | Blue-throated Barbet |  | × | R | LC | Schedule IV |
| *Psilopogon virens* (Boddaert, 1783) | Great Barbet | × | × | R/AM | LC | Schedule IV |
| **Falconiformes (2)** | | | | | | |
| **Falconidae (2)** | | | | | | |
| *Falco naumanni* Fleischer, 1818 | Lesser Kestrel |  | × | PV | LC | Schedule IV |
| *Falco tinnunculus* Linnaeus, 1758 | Common Kestrel | × | × | WV/R | LC | Schedule IV |
| **Psittaciformes (1)** | | | | | | |
| **Psittaculidae (1)** | | | | | | |
| *Psittacula himalayana* (Lesson, 1832) | Slaty-headed Parakeet | × |  | R | LC | Schedule IV |

| **Scientific Name** | **Common Name** | **Sighted at** | | **Migratory Status** | **IUCN status** | **IWPA status** |
| --- | --- | --- | --- | --- | --- | --- |
|  |  | **Nainital District** | **Mussoorie District** |  |  |  |
| **Passeriformes (83)** | | | | | | |
| **Campephagidae (2)** | | | | | | |
| *Pericrocotus ethologus* Bangs & J.C. Phillips, 1914 | Long-tailed Minivet | × |  | WV/R/AM | LC | Schedule IV |
| *Pericrocotus flammeus* (J.R. Forster, 1781) | Scarlet Minivet | × |  | R/AM | LC | Schedule IV |
| **Vireonidae (1)** | | | | | | |
| *Pteruthius ripleyi* Biswas, 1960 | Himalayan Shrike-babbler | × | × | R/AM | LC | Schedule IV |
| **Oriolidae (1)** | | | | | | |
| *Oriolus traillii* (Vigors, 1832) | Maroon Oriole | × |  | R/AM | LC | Schedule IV |
| **Dicruridae (4)** | | | | | | |
| *Dicrurus annectens* (Hodgson, 1836) | Crow-billed Drongo | × |  | SV | LC | Schedule IV |
| *Dicrurus caerulescens* (Linnaeus, 1758) | White-bellied Drongo | × |  | WV/AM | LC | Schedule IV |
| *Dicrurus leucophaeus* Vieillot, 1817 | Ashy Drongo | × |  | SV/R | LC | Schedule IV |
| *Dicrurus macrocercus* Vieillot, 1817 | Black Drongo | × |  | R/AM | LC | Schedule IV |
| **Rhipiduridae (1)** | | | | | | |
| *Rhipidura albicollis* (Vieillot, 1818) | White-throated Fantail | × | × | R/AM | LC | Schedule IV |
| **Laniidae (1)** | | | | | | |
| *Lanius tephronotus* (Vigors, 1831) | Grey-backed Shrike | × | × | WV | LC | Schedule IV |
| **Corvidae (5)** | | | | | | |
| *Corvus macrorynchos* Wagler, 1827 | Large-billed Crow | × |  | R | LC | Schedule IV |
| *Dendrocitta formosae* Swinhoe, 1863 | Grey Treepie | × |  | R | LC | Schedule IV |
| *Garrulus glandarius* (Linnaeus, 1758) | Eurasian Jay | × |  | R | LC | Schedule IV |
| *Garrulus lanceolatus* Vigors, 1830 | Black-headed Jay | × |  | R/AM | LC | Schedule IV |
| *Urocissa erythroryncha* (Boddaert, 1783) | Red-billed Blue Magpie | × |  | R/AM | LC | Schedule IV |
| **Dicaediae (1)** | | | | | | |
| *Dicaeum ignipectus* (Blyth, 1843) | Fire-breasted Flowerpecker | × |  | R/AM | LC | Schedule IV |
| **Nectariniidae (1)** | | | | | | |
| *Aethopyga siparaja* (Raffles, 1822) | Crimson Sunbird | × |  | R/AM | LC | Schedule IV |
| **Prunellidae (1)** | | | | | | |
| *Prunella himalayana* (Blyth, 1842) | Himalayan Accentor | × | × | WV | LC | Schedule IV |
| **Passeridae (2)** | | | | | | |
| *Passer cinnamomeus* (Gould, 1836) | Russet Sparrow |  | × | R/AM | LC | Schedule IV |
| *Passer domesticus* (Linnaeus, 1758) | House Sparrow | × | × | R/AM | LC | Schedule IV |
| **Motacillidae (2)** | | | | | | |
| *Motacilla alba* Linnaeus, 1758 | White Wagtail | × |  | WV | LC | Schedule IV |
| *Motacilla cinerea* Tunstall, 1771 | Grey Wagtail | × |  | WV/AM | LC | Schedule IV |
| **Fringillidae (5)** | | | | | | |
| *Carpodacus rodochroa* (Vigors, 1831) | Pink-browed Rosefinch |  | × | WV | LC | Schedule IV |
| *Chloris spinoides* (Vigors, 1831) | Yellow-breasted Greenfinch | × |  | WV/R | LC | Schedule IV |
| *Erythrina erythrina* (Pallas, 1770) | Common Rosefinch |  | × | WV | LC | Schedule IV |
| *Mycerobas carnipes* (Hodgson, 1836) | White-winged Grosbeak |  | × | PV/AM | LC | Schedule IV |
| *Pyrrhula nipalensis* Hodgson, 1836 | Brown Bullfinch |  | × | AM/WV | L | Schedule IV |
| **Emberizidae (2)** | | | | | | |
| *Emberiza cia* Linnaeus, 1766 | Rock Bunting | × | × | WV | LC | Schedule IV |
| *Emberiza stewarti* (Blyth, 1854) | White-capped Bunting | × |  | WV | LC | Schedule IV |
| **Stenostiridae (1)** | | | | | | |
| *Culicicapa ceylonensis* (Swainson, 1820) | Grey-headed Canary-flycatcher | × |  | R/AM/WV | LC | Schedule IV |
| **Paridae (5)** | | | | | | |
| *Machlolophus xanthogenys* (Vigors, 1831) | Himalayan Black-lored Tit | × | × | R/AM | LC | Schedule IV |
| *Parus cinereus* Vieillot, 1818 | Cinerous Tit |  | × | R | LC | Schedule IV |
| *Parus monticolus* Vigors, 1831 | Green-backed Tit | × | × | R | LC | Schedule IV |
| *Periparus ater* (Linnaeus, 1758) | Coal Tit | × | × | R/AM | LC | Schedule IV |
| *Sylviparus modestus* E. Burton, 1836 | Yellow-browed Tit | × | × | R/AM | LC | Schedule IV |
| **Cisticolidae (1)** | | | | | | |
| *Prinia crinigera* Hodgson, 1836 | Striated Prinia | × |  | R/AM | LC | Schedule IV |
| **Hirundinidae (1)** | | | | | | |
| *Delichon nipalense* F. Moore, 1854 | Nepal House Martin | × |  | R/AM | LC | Schedule IV |
| **Pycnonotidae (4)** | | | | | | |
| *Hypsipetes leucocephalus* (J.F. Gmelin, 1789) | Himalayan Black Bulbul | × | × | R/AM | LC | Schedule IV |
| *Ixos mcclellandii* (Horsfield, 1840) | Mountain Bulbul | × |  | R/AM | LC | Schedule IV |
| *Pycnonotus cafer* (Linnaeus, 1766) | Red-vented Bulbul | × | × | R | LC | Schedule IV |
| *Pycnonotus leucogenis* (J.E. Gray, 1835 ) | Himalayan Bulbul | × | × | R | LC | Schedule IV |
| **Phylloscopidae (6)** | | | | | | |
| *Abrornis chloronotus* J.E. & G.R. Gray, 1847 | Lemon-rumped Warbler | × | × | WV | LC | Schedule IV |
| *Abrornis maculipennis* Blyth, 1867 | Ashy-throated Warbler | × | × | AM/WV | LC | Schedule IV |
| *Abrornis pulcher* (Blyth, 1845) | Buff-barred Warbler | × | × | AM/WV | LC | Schedule IV |
| *Phylloscopus griseolus* Blyth, 1847 | Sulphur-bellied Warbler | × |  | SV | LC | Schedule IV |
| *Seicercus nitidus* (Blyth, 1843) | Green Leaf Warbler | × | × | PV/SV | LC | Schedule IV |
| *Seicercus xanthoschistos* (J.E. & G.R. Gray, 1847) | Grey-hooded Leaf Warbler | × | × | R/AM | LC | Schedule IV |

| **Scientific Name** | **Common Name** | **Sighted at** | | **Migratory Status** | **IUCN status** | **IWPA status** |
| --- | --- | --- | --- | --- | --- | --- |
|  |  | **Nainital District** | **Mussoorie District** |  |  |  |
| **Scotocercidae (2)** | | | | | | |
| *Abroscopus schisticeps* (J.E. & G.R. Gray, 1847) | Black-faced Warbler | × | × | R/AM | LC | Schedule IV |
| *Horornis fortipes* Hodgson, 1845 | Brownish-flanked Bush Warbler |  | × | WV/AM | LC | Schedule IV |
| **Aegithalidae (1)** | | | | | | |
| *Aegithalos concinnus* (Gould, 1855) | Black-throated Tit | × | × | R | LC | Schedule IV |
| **Zosteropidae (3)** | | | | | | |
| *Yuhina flavicollis* Hodgson, 1836 | Whiskered Yuhina | × | × | R/AM | LC | Schedule IV |
| *Yuhina gularis* Hodgson, 1836 | Stripe-throated Yuhina |  | × | R/AM | LC | Schedule IV |
| *Zosterops palpebrosus* (Temminck, 1824) | Oriental White-eye | × | × | R | LC | Schedule IV |
| **Timaliidae (2)** | | | | | | |
| *Cyanoderma pyrrhops* (Blyth, 1844) | Black-chinned Babbler |  | × | R | LC | Schedule IV |
| *Erythrogenys erythrogenys* (Vigors, 1831) | Rusty-cheeked Scimitar Babbler |  | × | R | LC | Schedule IV |
| **Leiothrichidae (4)** | | | | | | |
| *Garrulax albogularis* (Gould, 1836) | White-throated Laughingthrush | × | × | R/AM | LC | Schedule IV |
| *Grammatoptila striata* (Vigors, 1831) | Striated Laughingthrush | × | × | R | LC | Schedule IV |
| *Heterophasia capistrata* (Vigors, 1831) | Rufous Sibia | × | × | R | LC | Schedule IV |
| *Trochalopteron lineatum* (Vigors, 1831) | Streaked Laughingthrush | × | × | R | LC | Schedule IV |
| **Certhiidae (1)** | | | | | | |
| *Certhia himalayana* Vigors, 1832 | Bar-tailed Treecreeper | × | × | R | LC | Schedule IV |
| **Sittidae (2)** | | | | | | |
| *Sitta himalayensis* Jardine & Selby, 1835 | White-tailed Nuthatch | × | × | R/AM | LC | Schedule IV |
| *Tichodroma muraria* (Linnaeus, 1766) | Wallcreeper |  | × | WV | LC | Schedule IV |
| **Troglodytidae (1)** | | | | | | |
| *Troglodytes troglodytes* (Linnaeus, 1758) | Eurasian Wren |  | × | WV | LC | Schedule IV |
| **Sturnidae (1)** | | | | | | |
| *Acridotheres tristis* (Linnaeus, 1766) | Common Myna | × | × | R/AM | LC | Schedule IV |
| **Muscicapidae (13)** | | | | | | |
| *Enicurus schistaceus* (Hodgson, 1836) | Slaty-backed Forktail | × |  | AM | LC | Schedule IV |
| *Eumyias thalassinus* (Swainson, 1838) | Verditer Flycatcher | × |  | SV/AM | LC | Schedule IV |
| *Ficedula superciliaris*(Jerdon, 1840) | Ultramarine Flycatcher | × |  | SV/AM | LC | Schedule IV |
| *Monticola rufiventris* (Jardine & Selby, 1833) | Rufous-bellied Rock Thrush | × | × | R/AM | LC | Schedule IV |
| *Muscicapa sibirica* J.F. Gmelin, 1789 | Dark-sided Flycatcher | × |  | SV/AM | LC | Schedule IV |
| *Myophonus caeruleus* (Scopoli, 1786) | Blue Whistling Thrush | × | × | R/AM | LC | Schedule IV |
| *Niltava sundara* Hodgson, 1837 | Rufous-bellied Niltava | × |  | R/AM | LC | Schedule IV |
| *Phoenicurus coeruleocephala* (Vigors, 1831) | Blue-capped Redstart | × | × | AM/WV | LC | Schedule IV |
| *Phoenicurus frontalis* (Vigors, 1831) | Blue-fronted Redstart | × | × | AM/WV | LC | Schedule IV |
| *Phoenicurus fuliginosa* (Vigors, 1831) | Plumbeous Water Redstart | × |  | R/AM | LC | Schedule IV |
| *Phoenicurus leucocephalus* (Vigors, 1831) | White-capped Water Redstart | × |  | AM/WV | LC | Schedule IV |
| *Saxicola ferreus* J.E. & G.R. Gray, 1847 | Grey Bushchat | × | × | R/AM | LC | Schedule IV |
| *Tarsiger rufilatus* (Hodgson, 1845) | Himalayan Bluetail | × | × | AM/WV | LC | Schedule IV |
| **Turdidae (thrushes) (6)** | | | | | | |
| *Turdus atrogularis* Jarocki, 1819 | Black-throated Thrush | × | × | WV | LC | Schedule IV |
| *Turdus boulboul* (Latham, 1790) | Grey-winged Blackbird | × | × | R/AM | LC | Schedule IV |
| *Turdus rubrocanus* J.E. & G.R. Gray, 1847 | Chestnut Thrush | × | × | AM/WV | LC | Schedule IV |
| *Turdus unicolor* Tickell, 1833 | Tickell’s Thrush | × |  | SV/AM | LC | Schedule IV |
| *Zoothera dauma* (Latham, 1790) | Scaly Thrush | × |  | AM | LC | Schedule IV |
| *Zoothera mollissima* (Blyth, 1842) | Plain-backed Thrush | × | × | WV/AM | LC | Schedule IV |

*Note*: These 116 species are those found in the probable altitudinal range of *O.superciliosa*, in the study area (appox. 1600 m and above). The rest of the 162 confirmed sightings/calls and other 8 unconfirmed species have been omitted. Taxonomy is based on Praveen et al (2016, 2018)^[1][2]^. The migratory status has been determined for the two districts using species range maps from del Hoyo et al (2020)^[3]^, Grimmett and Inskipp (2014)^[4]^ and records from eBird.

Key

Migratory status: R = Resident; AM = Altitudinal Migrant; SV = Summer Visitor; WV = Winter Visitor; PV = Passage (Autumn and/or Spring) Visitor IUCN status: LC = Least Concern; VU = Vulnerable; NT = Near Threatened

^[1]^ Praveen, J., Jayapal, R., Pittie, A. (2016). A checklist of the birds of India. *Indian BIRDS*, *11*(5 & 6), 113-172A. <https://www.researchgate.net/publication/305307630_A_checklist_of_the_birds_of_India> Last accessed: 10^th^ July 2020

^[2]^ Praveen, J., Jayapal, R., & Pittie, A. (2018). Taxonomic updates to the checklist of birds of India and the South Asian region. *Indian BIRDS*, *14*(2), 37-42. <http://indianbirds.in/pdfs/IB_14_2_PraveenETAL_TaxonomicUpdates2018.pdf> Last accessed: 10^th^ July 2020

^[3]^del Hoyo, J., Elliott, A., Sargatal, J., Christie, D. A., & de Juana, E. (Eds) (2020). *Birds of the World*. Cornell Lab of Ornithology. [https://doi.org/10.2173/bow](https://doi.org/10.2173/bow.bfbwar1.01)

^[4]^ Grimmett, R., Inskipp, C., & Inskipp, T. (2014). *Birds of the Indian Subcontinent*. London, UK:	Bloomsbury Publishing.
